# ‘broadSeq’, a benchmarking tool to contrast differential gene expression methods and identify when genomic neighbourhood matters

**DOI:** 10.64898/2026.10.03.756406

**Authors:** Rishi Das Roy, Outi K. Hallikas, Satu H. Kuure, Jukka Jernvall

## Abstract

Several tools are available to detect differential gene expression. The predominant approach of different statistical methods is to consider changes in gene expression independent from their genomic context. Previously, DELocal was introduced that takes into consideration gene expression dynamics in the genomic neighbourhood. Whereas DELocal was shown to identify developmentally regulated genes in developing mouse tooth, its performance in other biological conditions remains undetermined. To benchmark DElocal together with other methods, we develop another tool ‘broadSeq’. It provides easy to use interface for methods like DELocal, deseq2, limma, edger, EBSeq and many more. Here we report benchmarking results on different data sets that comprise wild type developing mouse organs from different embryonic time points, and also human cancer data. The examined datasets are from the heart, kidney, liver, brain, forebrain, and hindbrain. We found that most methods produce relatively comparable lists of differentially expressed genes. Nevertheless, DELocal identifies partially distinct sets of genes compared to the other methods. Evaluating the identified genes using gene and disease ontology terms show that the uniquely detected genes by DELocal are biologically relevant and also highly expressed compared to the genes identified by the other methods. Because consistently highly expressed genes show relatively subtle fold changes, the neighbourhood concept of DELocal appears to correct for the difficulty in detecting these differential expression of highly expressed genes. Finally, we provide the tools as combined bioconductor packages, allowing researchers to explore and decide the best methods for their data and questions.

## Introduction

One of the many ways to understand biology at molecular level is precise measurement of gene expression, a task whose technological underpinnings have shifted dramatically from continuous microarray data to count-based sequencing. High-throughput sequencing has made it possible to simultaneously interrogate thousands of genes, yet this power comes with a steep computational cost: an overwhelming proliferation of tools for quality control, analysis, and detection of differentially expressed (DE) genes. These methods differ substantially in their statistical machinery, parametric versus non-parametric models, and different strategies for normalization, dispersion estimation, and multiple-testing correction (Chen et al. 2025, Leng et al. 2013, Love et al. 2014, Ritchie et al. 2015, Tarazona et al. 2011). Many of them gain their power by borrowing information across genes, shrinking gene-wise variance or dispersion estimates toward a genome-wide trend. That pooling, however, treats genes as exchangeable and serves only to stabilize nuisance parameters: the quantity being tested remains the change in each gene’s own expression between conditions. Perhaps partly as a consequence of this shared premise, no method has emerged as universally optimal. As demonstrated across spike-in and simulated datasets, the top-performing method varies between datasets, making it clear that a one-size-fits-all solution does not exist (Jabato et al. 2021). On the other hand, some studies show that different methods start to converge as sample size increase (Schurch et al. 2016, Soneson & Delorenzi 2013).

Nevertheless, the underlying principles how different methods work do matter as much as its statistical implementation. For example, DELocal, departs from the gene-in-isolation premise by measuring the expression change of a gene *relative to the expression of its neighbouring genes* (Das Roy et al. 2021). Its underlying neighbourhood concept of gene regulation assumes that the expression dynamics of all genes within a genomic region must be taken into account when determining whether a gene is differentially expressed. Because the reference point is local rather than global, a gene can be called differentially expressed when it deviates from its genomic neighbourhood even if its absolute fold change is modest, a regime in which fold-change-centric methods are comparatively insensitive. In practice, this shift in reference point has been shown to help: DELocal identified biologically well-validated DE genes with comparatively superior performance in developing tooth data of the mouse (Das Roy et al. 2021). That evidence, however, comes from a single organ and a single experimental design. Whereas genomic neighbourhood may be expected to be relevant for a developing organ with relatively fixed set of expressed genes, the applicability of DELocal across diverse transcriptomic datasets from different organs and experiments remain uncharacterized. Establishing whether the neighbourhood concept can be generalized requires that DELocal be run and compared against different methods on many datasets and contrasts.

The incompatible input-output formats across tools — even for ostensibly identical outputs such as lists of differentially expressed genes — make cross-method comparison laborious at best and error-prone at worst (Figure 1A). The stakes of method selection are not trivial. Choosing an inappropriate DE method for a given dataset risks both false discoveries and missed biology. Despite this, the field has lacked a systematic, accessible framework to compare these methods — a gap with direct consequences for scientific conclusions. Existing graphical user interface (GUI) solutions are not architected to incorporate new methods, rendering them progressively obsolete. These challenges are compounded by severe usability barriers. The few GUI alternatives cover only a narrow subset of quality-checking and DE methods, producing correspondingly limited results Table S1, Table S2, and are hard to integrate with advanced workflows or add new methods (Dimitrov & Gu 2020, Haering & Habermann 2021, Helmy et al. 2021, Su et al. 2019, Sundararajan et al. 2019). At least one study covers a large number of methods and shares their code, but it lacks documentation preventing its usage (Baik et al. 2020). RNAseqR (Chao et al. 2019) which covers multiple DE methods, is not anymore available in latest Bioconductor release 3.23 (Gentleman et al. 2004), underscoring the brief availability of some of the tools. The compcodeR provides feature to integrate user defined functions but its performance metrics are designed for simulated data and when true differential expression status for each gene or gold standard data is already known (Soneson 2014). Yet another method, ExpHunterSuite (Jabato et al. 2021) is available, but only for few methods. Taken together, testing methods such as DELocal that uses different concepts for DE detection is hampered by the lack of frameworks and tools.

**Figure 1:**
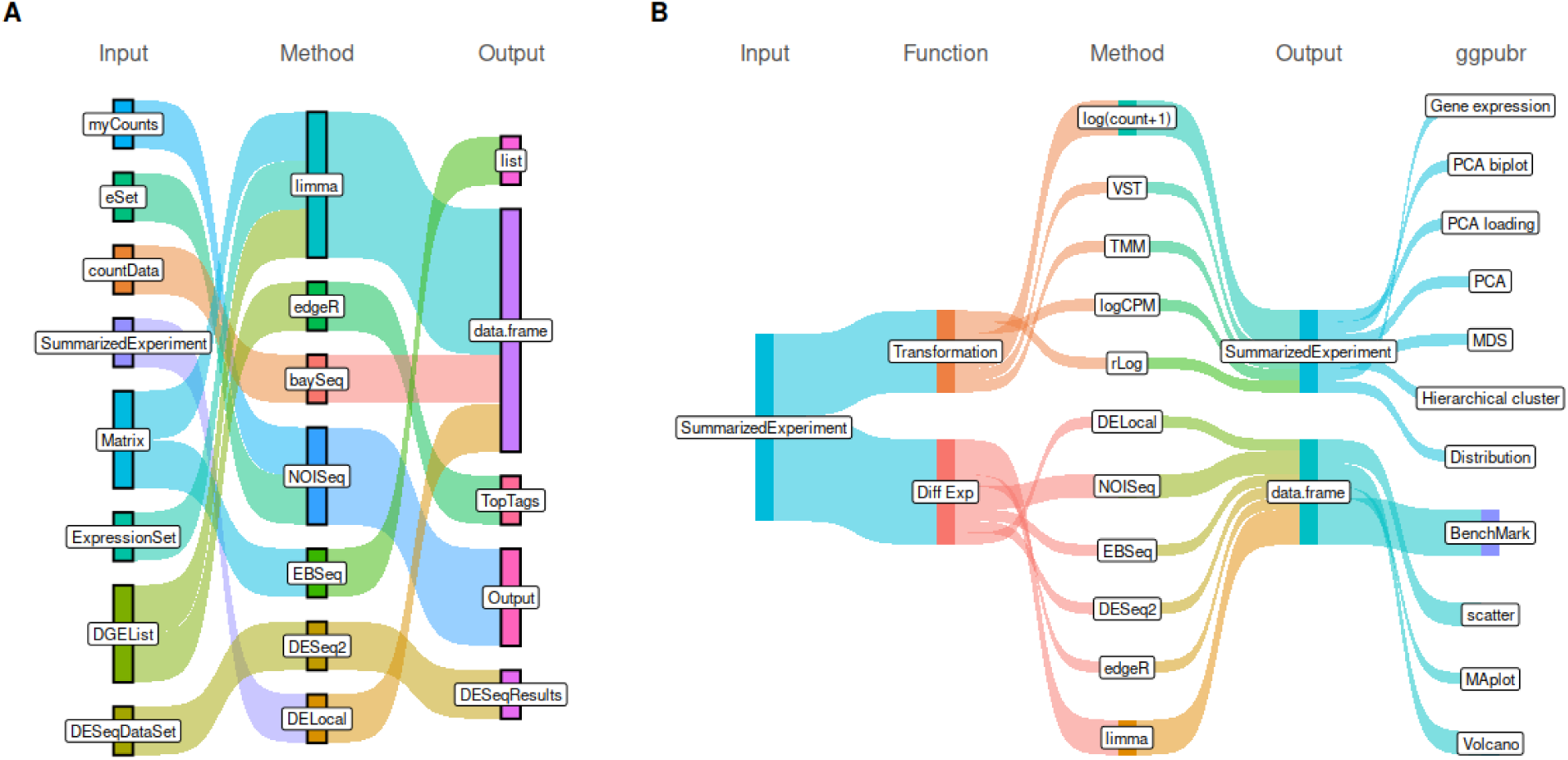
broadSeq streamlines the use of multiple RNA-seq tools. (A) Multiple input-output data formats are used by different tools. (B) broadSeq provides a simple interface to use these tools with only two different data formats: SummarizedExperiment and data.frame. These two methods are also compatible with a wide variety of approaches for graphical visualisation.

To address the absence of ready-to-use tools capable of systematically comparing different DE methods on user-provided data, here we present broadSeq, an R package whose input and output formats adhere to widely accepted standards. broadSeq internally handles data format conversion, enabling users to benchmark a broad range of DE methods without learning method-specific data structures (Figure 1B). Crucially, broadSeq is designed with extensibility at its core — developers and advanced users can introduce new DE methods into the comparison framework by implementing standardized wrapper functions within the broadSeq environment, ensuring the package remains relevant as the field evolves. Importantly, to overcome the absence of a ‘gold standard’ for evaluation, we leverage Gene Ontology (GO) as well as Disease Ontology (DO) term enrichments as proxies for biological relevance. That is, methods that recover more coherent, functionally interpretable GO and DO gene sets are considered superior. Here we apply broadSeq to benchmark DELocal against other methods on both mouse and human transcriptomic datasets. We demonstrate that DELocal not only identifies a unique set of differentially expressed genes but also recovers many biologically relevant GO and DO terms, substantiating its performance beyond the mouse developing tooth context in where it was originally characterized. Overall, broadSeq integrating GO/DO-based evaluation strategy provides a practical, extensible solution for rigorous DE method benchmarking across diverse transcriptomic datasets.

## Results

### broadSeq

A fundamental obstacle to multi-method DE analysis is that each method imposes its own input data structure, forcing researchers to redundantly reformat the same underlying data. This is not merely inconvenient, but maintaining parallel representations of the same dataset across multiple formats introduces a tangible risk of data management errors and inconsistencies within a project. The problem is further compounded by the fact that gene expression data is not simply a count matrix, as a complete representation must also encode sample-specific metadata as well as gene-specific annotations such as alternative symbols, biotypes, genomic coordinates, read length, and normalized abundance values.

The broadSeq resolves this by accepting input exclusively in the SummarizedExperiment format, a widely adopted Bioconductor standard capable of simultaneously holding raw counts, gene length, normalized values, sample metadata, and gene-level annotations in a single coherent object (Morgan et al. 2025). Rather than requiring users to learn and maintain method-specific data structures, broadSeq internally converts this unified representation into whatever format each downstream method requires. This design eliminates redundant data copies and substantially reduces the burden of reformating the data. From this single entry point, broadSeq offers a consistent interface for a broad range of analytical tasks, including expression scaling and normalization, exploratory visualization, sample clustering, and differential expression analysis across multiple methods.

### Normalization/Transformation

In RNA-seq analysis, gene expression is quantified by counting the number of sequencing reads aligning to each gene (Wang et al. 2009). These raw counts, however, are not directly comparable across samples without normalization, which is essential both for visualization and for compatibility with downstream analytical tools (Evans et al. 2018). Normalization strategies differ substantially in their assumptions and dependencies. Length-dependent units, RPKM (reads per kilobase of transcript per million mapped reads), FPKM (fragments per kilobase of transcript per million mapped reads), and TPM (transcripts per million), require isoform-level length information that is most accurately obtained at the alignment stage. Modern pseudo-aligners such as salmon (Patro et al. 2017a) and kallisto (Bray et al. 2016) simultaneously give as output raw counts, gene lengths, and TPM abundance estimates, and their output can be imported directly into the SummarizedExperiment format via the tximport package. For count-based normalization, edgeR provides CPM (counts per million) and TMM (trimmed mean of M-values), while DESeq2 offers rLog, VST (variance stabilizing transformation), and NTD (normalized transformation) transformations — yet these two packages accept mutually incompatible input formats, forcing users to maintain separate data objects for the same experiment. broadSeq eliminates this fragmentation: it provides a unified interface to all of these normalization and transformation methods and, critically, stores every transformation within the same SummarizedExperiment object, preserving a single, consistent data structure throughout the analysis. To demonstrate these capabilities, we applied broadSeq to RNA-seq data from developing tooth tissue across three rodent species, mouse, rat, and vole, spanning four developmental stages: bud, cap, late cap, and bell stage (Figure S1A) The effect of different normalization strategies can be assessed using broadSeq’s built-in boxplot functions, visualized either across samples (Figure S1B) or for individual genes of interest (Figure S2). Expression of multiple genes across normalization units, together with gene-level metadata, can be simultaneously displayed using dedicated built-in functions (Figure S3).

### Experiment-level quality control

Before differential expression analysis, it is essential to verify that the data contain sufficient signal to distinguish biologically distinct sample groups, a step commonly referred to as experiment-level quality control. In RNA-seq analysis, this is typically achieved by visualizing inter-sample relationships using dimensionality reduction and clustering methods, including hierarchical clustering, multidimensional scaling (MDS), and principal component analysis (PCA). Effective quality control further requires the ability to map sample metadata, such as treatment condition, developmental stage, or species onto these visualizations (using symbols such as shape, color, or size), and ideally to overlay gene-level annotations as well. Existing RNA-seq packages, however, neither natively support SummarizedExperiment objects nor offer more than one or two clustering methods within a single framework Table S1, Table S2, forcing users to write bespoke, non-reproducible code for each combination of scaling and clustering approach. broadSeq directly addresses this limitation by supporting any combination of the normalization and transformation methods described above as input to any available clustering method. Hierarchical clustering is implemented with integrated heatmap visualization, enabling simultaneous display of gene annotations alongside sample groupings (Figure 2A). MDS plots capture nonlinear inter-sample structure and are provided for rapid global overview (Figure 2B). PCA offers the most analytically rich feature set: beyond sample clustering, broadSeq exposes gene loadings on each principal component, which can be visualized as biplots or ranked barplots (Figure 2C, D). This is particularly valuable in complex experimental designs where gene expression is a function of more than two factors, necessitating systematic exploration of multiple PC combinations to fully characterize the sources of variation. In the rodent tooth dataset, for instance, PC3 captures variation attributable to developmental stage, a signal that is not recoverable through hierarchical clustering, or MDS alone. [The ability to interrogate gene–PC relationships through a simple, consistent interface is, to our knowledge, not available in any other RNA-seq package.]

**Figure 2:**
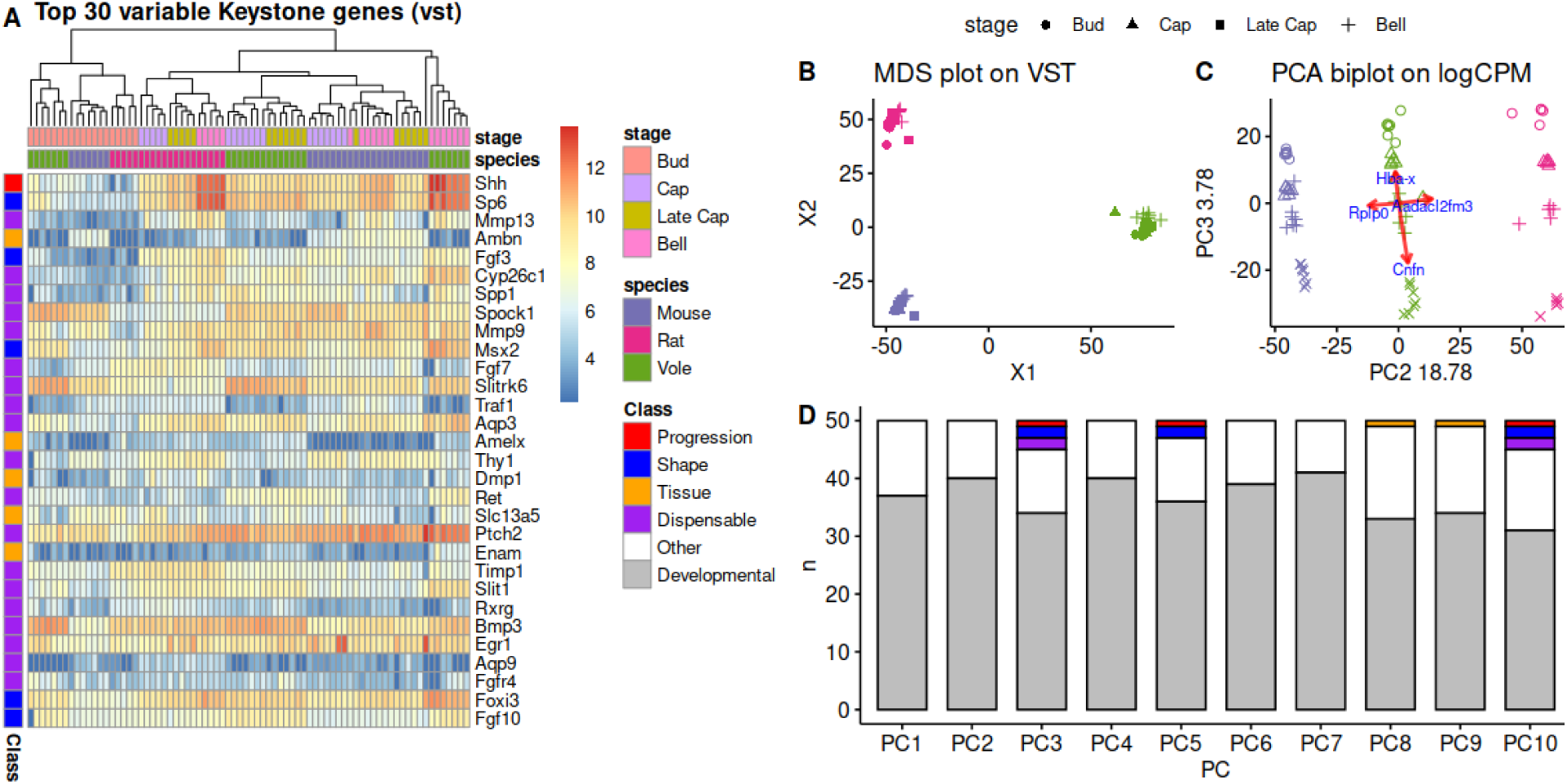
broadSeq offers different clustering methods to illuminate various features within the data. (A) Clustered heatmap with Keystone genes important for tooth development. (B) Multidimensional Scaling Plot with vst separates the samples by species. (C) PCA biplot with LogCPM explain variation due to species and developmental stages. (D) Top 50 genes loaded in the first 10 principal components coloured by different classes of Keystone genes.

### Differential expression analysis

broadSeq implements a comprehensive suite of DE methods which includes limma(Ritchie et al. 2015), DESeq2(Love et al. 2014), edgeR(Chen et al. 2025), EBSeq(Leng et al. 2013), NOISeq(Tarazona et al. 2011) and DELocal(Das Roy et al. 2021). Its API-oriented architecture allows users to introduce additional methods as wrapper functions, ensuring the framework remains extensible as new approaches emerge. Despite the abundance of available DE methods, there is no consensus on which method should be preferred. The choice is non-trivial as methods differ fundamentally in their statistical or biological assumptions (e.g., parametric versus non-parametric, using neighbourhood expression data) and in how they handle dispersion estimation, normalization, and multiple testing.

In practice, these differences force researchers to run each method independently, reformatting input data into incompatible structures for every tool, a workflow that is error-prone, time-consuming, and rarely reproducible (Figure 1A). The absence of a unified evaluation framework is therefore not merely an inconvenience; it is a structural barrier to method benchmarking in applied research.

For DE analyses, broadSeq provides a single, consistent interface to eight DE methods, including two variants each of limma and edgeR (two additional packages were excluded due to instability). From this interface, results across all methods are returned in a common format, enabling straightforward cross-method comparison without any additional data wrangling. Results can be visualized using volcano plots, MA plots, and scatter plots — a combination not fully available in any individual package or existing meta-tool Table S1, Table S2.

### Method Comparison

To benchmark DE methods across diverse biological contexts, we utilized a public RNA-seq dataset of developing mouse organs — heart, liver, brain, forebrain, hindbrain, and kidney (Figure 3A) (Cardoso-Moreira et al. 2019). This dataset is well-suited for benchmarking because it encompasses two sources of biological variation: organ identity and developmental trajectory. These sources of variation are distributed across many principal components (Figure S4A–B), and their underlying structure across developmental trajectories and stages is summarized in (Figure 3B).

**Figure 3:**
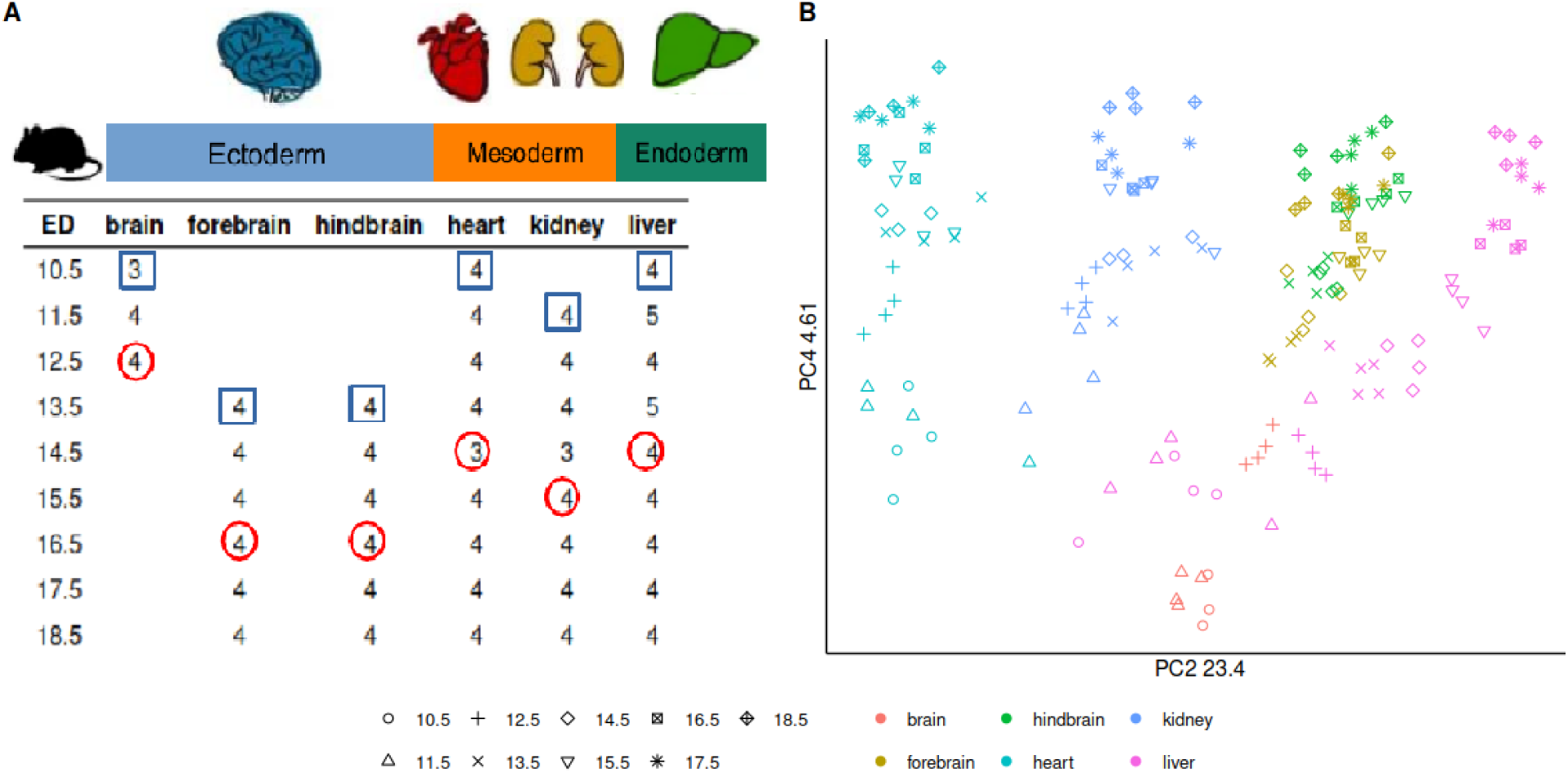
Transcriptomic data of mouse from multiple embryonic developmental stages of six different organs are used to evaluate the methods. (A) The numbers of replicates for each biological condition. Up-regulated genes are identified for two different contrasts i) one organ relative to all others and ii) at mid-development (red circles) relative to the earliest timepoint (blue square) (B) PCA plot of top 500 variable gene captures the gene expression pattern required to identify both organ specificity and developmental trajectory. Figure layout follows Cardoso-Moreira et al. (2019).

For each organ, two biologically meaningful contrasts were defined and only up-regulated genes were evaluated: (i) a given organ compared to all remaining organs, capturing organ-specific expression, and (ii) a middle developmental timepoint *t_m_* compared to the earliest timepoint *t*_0_, capturing developmental progression. For liver, *t_m_* and *t*_0_ correspond to embryonic days 14.5 and 10.5, respectively (Figure 3A). These contrasts were chosen deliberately, as organ-specificity and temporal induction represent distinct modes of differential regulation, and a method that performs well across both provides stronger evidence of general utility.

Significant genes were selected for each method using either adjusted *p*-values or FDR (false discovery rate), depending on what each method reports. Genes showing significant upregulation were then ranked by their corresponding fold-change values to enable cross-method comparison on a common scale. Here DELocal, based on the neighbourhood concept, is an exception: it reports relative log fold-change values compared to neighbouring genes, and these relative values are used here for ranking. Ranking is critical to this evaluation because it moves beyond binary significance calls, as two methods may agree on the set of significant genes yet disagree substantially on their relative importance, and it is this ordering that ultimately determines which genes are prioritized for biological follow-up.

### Disparity/Similarity

To quantify the degree of agreement between different methods of differential gene expression analysis, we selected varying numbers of top-ranked genes (10, 20, 50, 100, 200, 500, and 1000) from each method for each contrast. As an initial visual assessment, Euler diagrams, a generalization of Venn diagrams, illustrate the overlap and uniqueness of each method’s output (Figure 4A). However, visual inspection alone is insufficient when the number of method combinations is large. For rigorous quantification, we computed the Jaccard index, which measures similarity of a method as the ratio of shared to total unique genes compared to the rest methods. Across all contrasts and all top-gene thresholds, DELocal consistently exhibited the lowest similarity to every other method (Figure S5), indicating that it identifies a distinct set of genes regardless of the stringency of selection.

**Figure 4:**
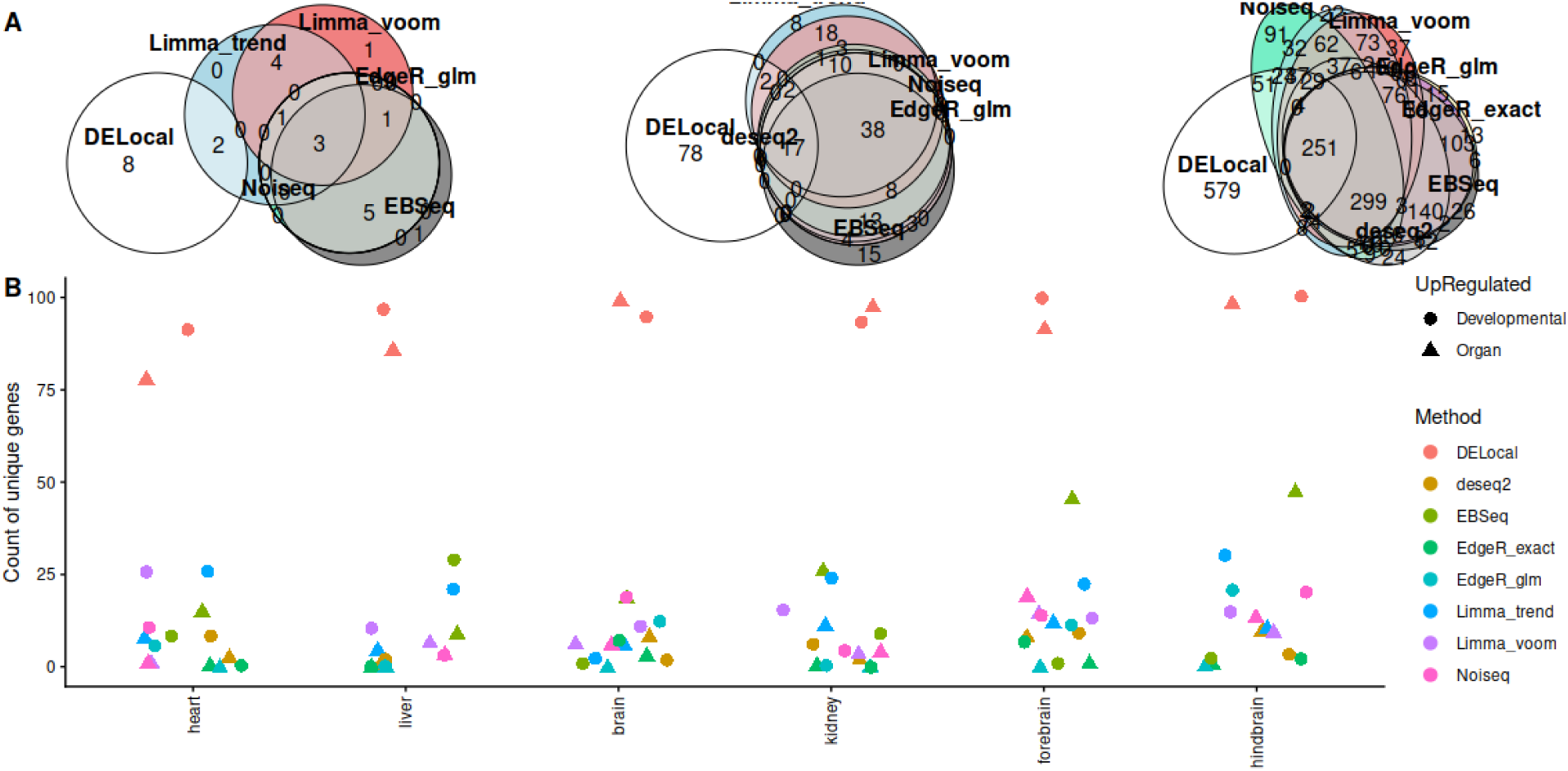
DELocal identifies maximum unique upregulated genes. A) Euler venn diagrams of top 10, 100 and 1000 upregulated genes in heart compared to rest of the organs. B) Top 100 upregulated genes identified by each method in two different contrasts; i) Organ vs rest and ii) middle developmental time t_m_ compared to t_0_

For more focused and tractable comparison, subsequent analyses concentrate on the top 100 ranked genes per contrast. Counting the unique genes reported exclusively by each method confirms the pattern observed above: the majority of methods show substantial mutual overlap, while DELocal stands as a consistent outlier, reporting a large number of highly ranked genes that are not identified as significant by any other method (Figure 4B). This is not a marginal difference; it reflects a qualitatively different detection strategy. The critical question this raises is not whether DELocal is different, but whether the unique genes detected using DELocal’s neighbourhood approach are biologically meaningful.

### Evaluation with gene ontology

To assess whether the genes uniquely identified by DELocal are biologically relevant rather than merely statistically distinct, we performed Gene Ontology (GO) enrichment analysis on the top-ranked gene sets from each method. The central question is whether DELocal and the other methods, despite their divergent gene lists, converge on biologically coherent and contextually appropriate GO terms for the developing organs under study, or whether DELocal’s unique genes reflect noise. GO enrichment was performed using the compareCluster function from the clusterProfiler package for “Biological Process” (BP), which enables simultaneous enrichment analysis across multiple gene lists (Xu et al. 2024, Yu et al. 2012). To reduce redundancy among enriched terms and improve interpretability, results were subsequently filtered using the simplify function with default parameters, which removes highly similar GO terms based on semantic similarity (Yu et al. 2012). We examined the results in the six different organs separately.

#### Heart

In the organ-versus-rest contrast, DELocal recovers GO terms consistent with those reported by other methods (Figure 5A) yet achieves this through 78 genes not identified by any other method (Figure 4B). The developmental trajectory contrast reveals an further differences. The GO terms “Extracellular matrix organization” and “connective tissue development”, which are relevant to development of heart (Del Monte-Nieto et al. 2020, Mendelson & Schoen 2006, Oliveira et al. 2025, Tallquist 2020), are found enriched in genes identified by DELocal (Figure 5B). Also, the GO terms “response to amino acid” and “cellular response to amino acid stimulus” are appropriate, as certain amino acids act as growth stimuli for cardiomyocyte proliferation (Li et al. 2025). Thus, DELocal appears to recovered several potentially meaningful GO terms which were not identified by other methods.

**Figure 5:**
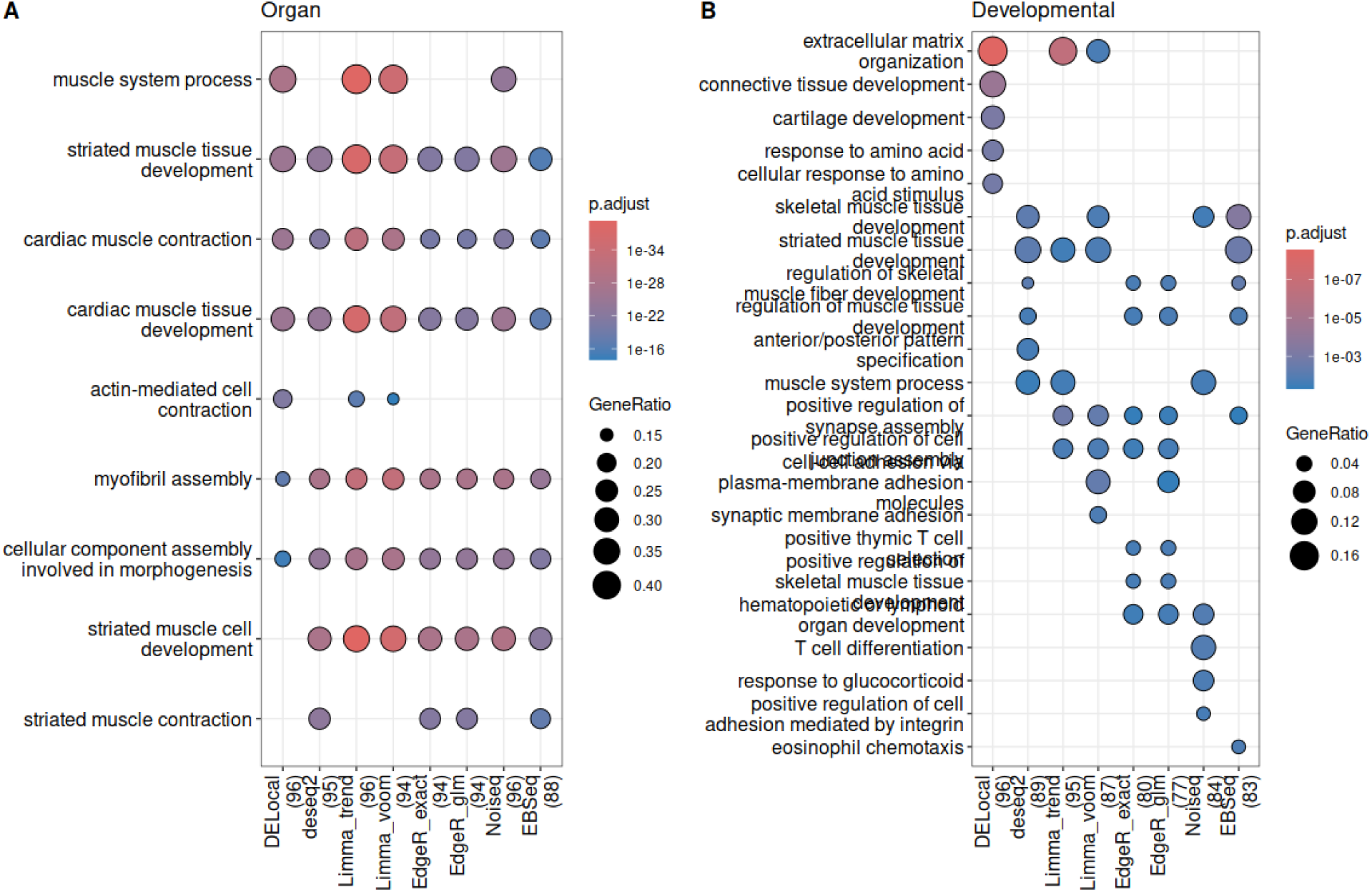
Over-representation analysis of GO terms in top 100 up regulated genes identified by different methods. The number of genes found in the annotation database shown next to each method. Up-regulated genes in (A) heart compared to the rest of organs and (B) mid-developmental time point (embryonic day 14.5) compared to earliest time point (embryonic day 10.5). In left, DELocal reports similar GO terms although 78 genes are unique.

#### Liver

In Gene Ontology (GO) enrichment analysis for liver samples, DELocal results were quite distinct from the other methods (Figure 6). Still, the GO terms retrieved by DELocal are relevant to liver function. Fetal liver is a major hematopoietic organ. During mid-gestation in mammals, the liver becomes the primary site of erythropoiesis (Ceccacci et al. 2023, McGrath & Palis 2008, Orkin & Zon 2008, Palis & Yoder 2001, Popescu et al. 2019). Thus, the processes described by GO terms like “erythrocyte development”, “erythrocyte homeostasis,” “hemoglobin biosynthetic process,” and “heme metabolic process” are directly active in the fetal liver (Figure 6A, B). In addition, liver stores and regulates the blood levels of many metal ions, most importantly iron (Anderson & Shah 2013). DELocal uniquely identifies this liver function in the GO terms “response to metal ion” and “cellular response to metal ion” (Figure 6A). These terms were recovered by DELocal but not by other methods. Therefore, even if DELocal does not recover all the GO terms retrieved by the other methods, it appears to provide biologically meaningful results.

**Figure 6:**
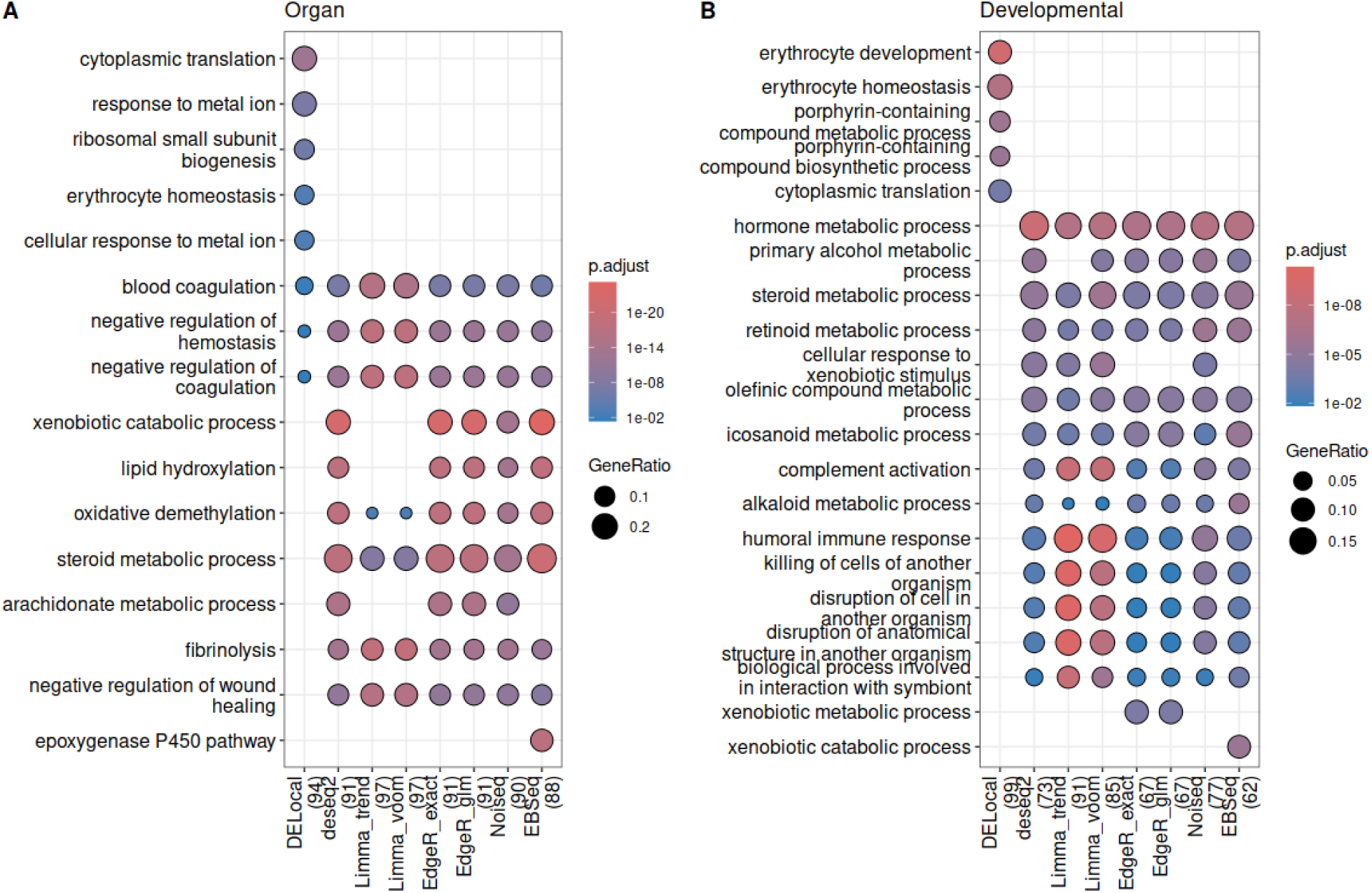
In both contrasts correspond to liver, DELocal stands out with relevant GO terms. (A) Up-regulated genes in Liver compared to the rest of organs (B) For developmentally upregulated genes, expression data from embryonic day 14.5 were compared to embryonic day 10.5

#### Brain, Forebrain, and Hindbrain

Across all contrasts involving brain, forebrain, and hindbrain, DELocal-reported genes are consistently enriched for distinct GO terms from those by all other methods, which converge on similar terms owing to the high overlap in their underlying gene lists (Figure S6, Figure S7, Figure S8). This pattern of DELocal divergence is reproducible across three related but anatomically distinct brain tissues, reinforcing that it is not a contrast-specific artifact. One exception warrants attention: in the brain-versus-rest contrast, the GO terms enriched in DELocal’s unique gene set are not strongly specific to brain function. This is likely an artifact of the comparison group composition, as forebrain and hindbrain are themselves included among the “rest” organs, which dilutes the brain-specific transcriptional signal and complicates the interpretation of any organ-versus-rest contrast for neuronal tissue. This is an inherent limitation of the contrast design rather than a failure of the method. For all remaining brain-related contrasts, including developmental trajectory comparisons within forebrain and hindbrain, DELocal uniquely recovers GO terms with direct neurobiological relevance, including terms associated with neuron differentiation, dendrite morphogenesis, synapse organization, and learning and memory (Figure 7). These are precisely the functional categories expected to be dynamically regulated during brain development, and their exclusive appearance in DELocal’s output makes a strong case that the identified genes are not statistical noise but biologically meaningful discoveries not captured by other methods.

**Figure 7:**
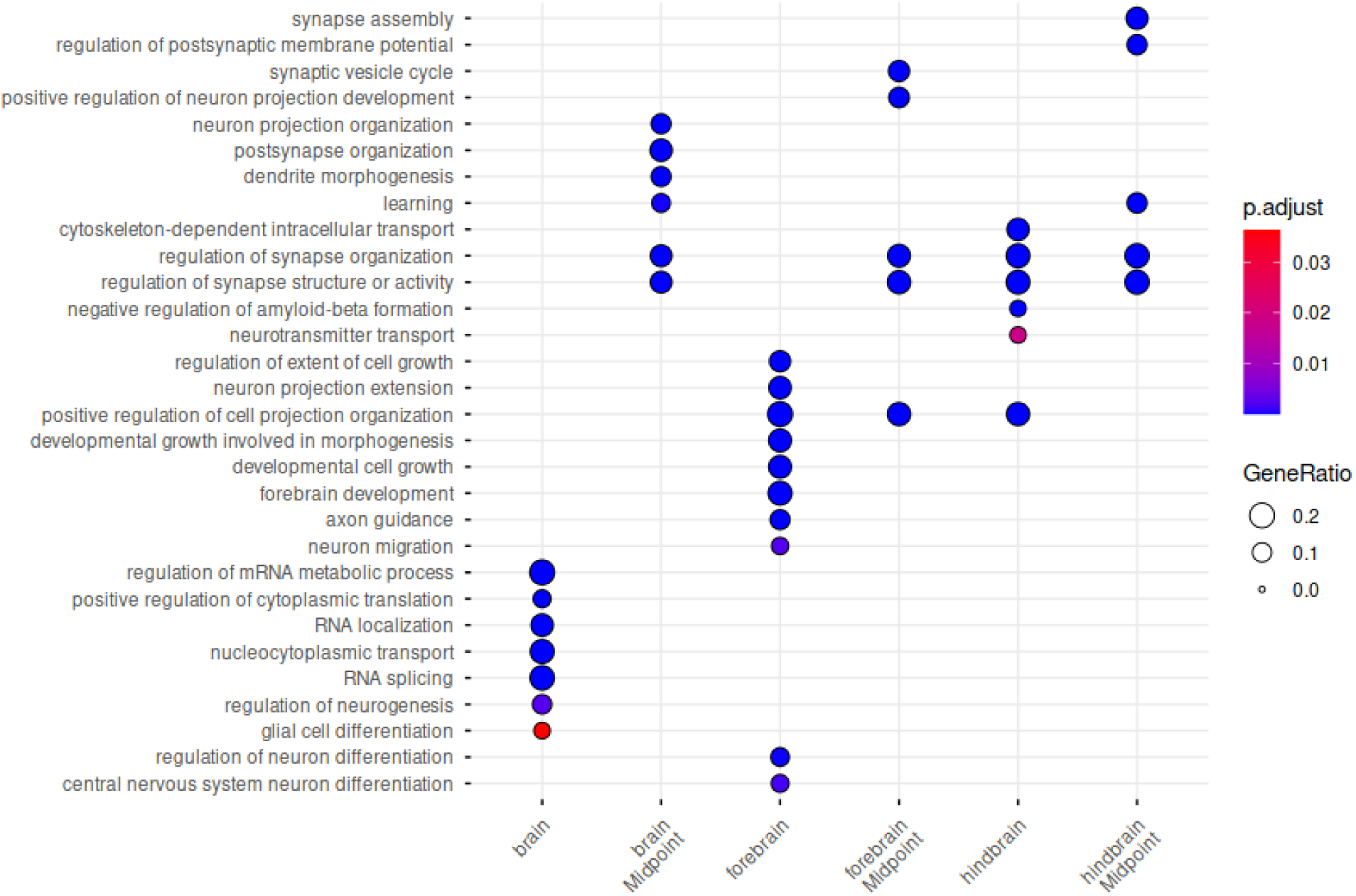
The unique GO-terms identified by DELocal in the contrasts of Brain, Forebrain, and Hindbrain are very relevant for the development of the brain. Except for brain-vs-rest, where GO terms are not relevant, which could be due to the presence of forebrain and hindbrain among the “rest” organs.

#### Kidney

For the organ-versus-rest contrast, genes up-regulated in developing kidney relative to all other organs are expectedly enriched with kidney-related GO terms across all the common methods. Consistent with the pattern observed in other organs, DELocal identifies unique but relevant GO terms not reported by any other method (Figure 8A), indicating that its neighbourhood-relative detection strategy captures genuine kidney-specific signal. The developmental trajectory contrast presents a more nuanced picture. At the midpoint of kidney development (E14.5)(Kurtzeborn et al. 2025, Kuure & Sariola 2020), DELocal’s enriched GO terms are less immediately recognizable than those of most other methods but reveal relevant metabolic processes related to core kidney functions, like amino acid breakdown and reabsorption (Figure 8B). Furthermore, when the full ranked output is examined, the top 20 significant GO terms from DELocal’s gene list include “kidney development” and “renal system development” (Figure S9) — terms that are directly pertinent to the contrast under study. Their absence from the main figure is a consequence of the clusterProfiler display limit imposed for readability, not an absence from the enrichment results. Taken together, the kidney development results suggest that DELocal identifies biologically coherent gene sets, even where superficial comparison of summary visualizations might suggest otherwise.

**Figure 8:**
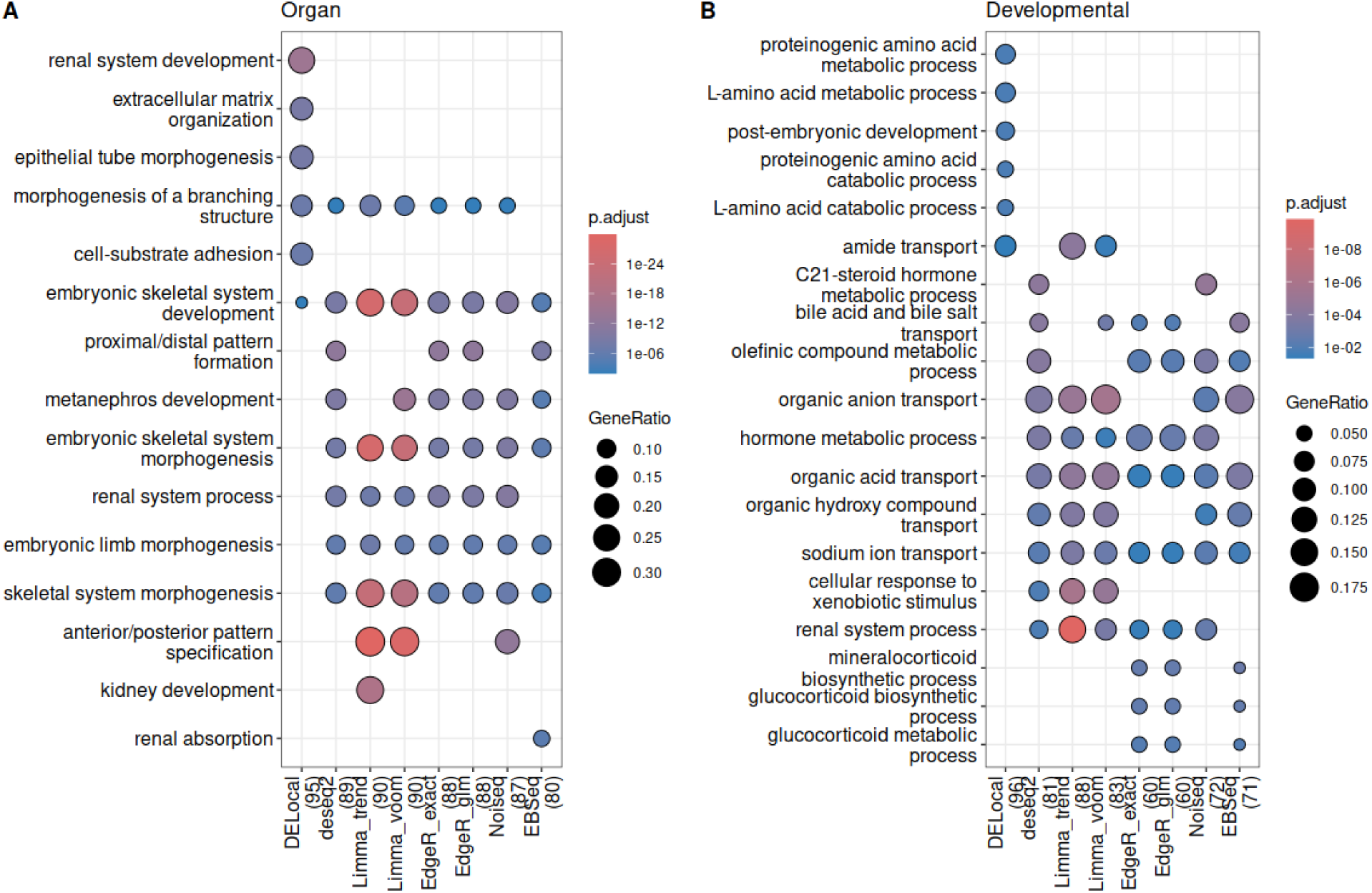
In both contrasts correspond to Kidney, (A & B) DELocal produces distinct GO terms which are associated with Kidney. However, none of the GO terms except ‘renal system process’ is immediately associated with Kidney.

#### Disease ontology

After applying broadSeq to evaluate different DE methods with gene ontology in mouse developing organs, next we compared and evaluate DE methods with disease ontology on a human kidney cancer data which is highly imbalanced with 6 normal and 124 diseased samples (Gadd et al. 2017) (Figure S10A). Probably due to the high-class imbalance between the control and cancer cases, only 4 methods in our study have reported top 100 DE genes (Figure 9A). Importantly, only DELocal detected nephroblastoma as a term and thus identified kidney cancer among other cancer related terms. Extended enriched disease ontology terms of DELocal identified genes also includes “kidney cancer” term (Figure S10B).

**Figure 9:**
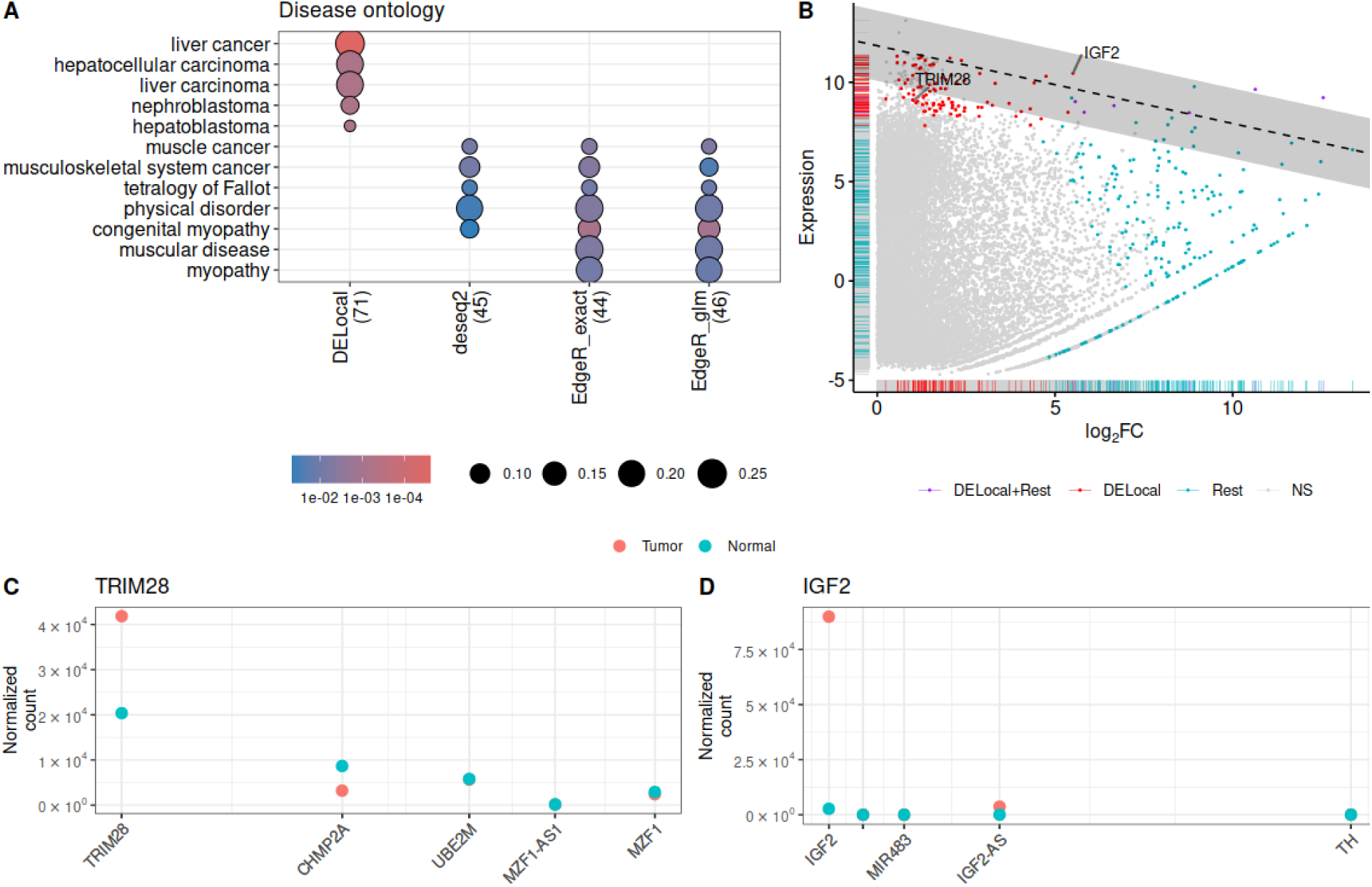
Up-regulated genes in human kidney cancer samples. (A) Evaluated with disease ontology (DO). The number of genes found in DO database shown next to the methods. (B) Scatter-plot of up-regulated genes shows DELocal identified genes are highly expressed and subtly differentially expressed (log fold change) compared to rest of the methods (measured by edgeR). TRIM28 and IGF2 which are not in DO database, have been recently found associated with Wilms tumor. (C) and (D) show their ‘median’ expression pattern compared to their neighbouring genes.

## Discussion & Conclusion

### broadSeq

The broadSeq provides an end-to-end pipeline for RNA-seq count data to perform comparative analysis, which is currently absent. It promotes the use of existing data structures and advanced analytical steps. The broadSeq generated publication-ready plots reduce the load on users and let them focus on their scientific objectives. Previous GUI based tools focus on user friendliness for naive researchers whereas API-oriented broadSeq is designed for experts who can build their own comparative workflow and choose best method to interpret their data. broadSeq provides alternatives not only for DE methods but also normalization and clustering methods. It promotes re-use of existing data structures and packages.

### DELocal

When examining how DELocal and its neighbourhood concept compares to the other methods, it can be observed that DELocal identified genes are highly expressed whereas their foldchange (*log*_2_) values tend to be lower than those detected by other methods (Figure 9B). In general, highly expressed genes tend to have lower fold changes (unless their expression plummets), resulting in lower rankings in typical DE lists. DELocal’s neighbourhood approach solve this problem by comparing the fold changes to changes in neighbouring genes. We note that simply selecting genes based on high expression levels would lead to false positives since many of these genes were not detected by either of the methods. The kidney cancer dataset was especially interesting as DELocal identified few genes that are still not associated with kidney cancer in the DO database. Performing a literature search on more recent studies to examine if any of the DELocal ranked genes are linked to kidney cancer resulted in two new cancer genes. TRIM28 (Hol et al. 2021, Murphy et al. 2023, Semenova et al. 2026, Wegert et al. 2024, 2025; Welter et al. 2023, Whitworth et al. 2024) and IGF2 (Bitaraf et al. 2022, Hong & Dong 2021, Jalal et al. 2024, Lim & Loh 2024, Wegert et al. 2025) are linked in recent studies to Wilms tumor, or nephroblastoma. Although these genes got significant *p-values* in the other methods, they had low rankings due to lower fold changes (TRIM28 = 1 and IGF2 = 5.5; Figure S11). In contrast, these two genes were highly ranked by DELocal. Overall it is plausible that many of the highly ranked DELocal genes may be true positive.

### Evaluation

The landscape of DE method benchmarking has been shaped by a recurring methodological tension: the need for a ground truth that real, noisy, biological data cannot provide. The earliest systematic comparison (Soneson & Delorenzi 2013) evaluated eleven methods both on synthetic counts, where the true positives were themselves synthetic, and real RNA-seq data, where in the absence of a verified gene list, shared DE genes across methods were used as a proxy for ground truth. While this study established the foundational framework for method comparison and demonstrated the utility of AUC-based (area under the curve) evaluation on simulated data, it did not address whether the identified genes — shared or unique — carried biological meaning. The question of relevance was left entirely open.

Subsequent studies inherited this limitation without resolving it. Seyednasrollah et al. (2015) applied hierarchical clustering and gene overlap counts to real mouse and human transcriptome data, but biological interpretation of method-specific genes remained absent. Tang et al. (2015) broadened the scope to multi-group designs, comparing twelve pipelines from nine R packages and recommending an iteratively normalized edgeR pipeline (DEGES/edgeR in TCC) for replicated data. Their real-data comparison, however, rested on the overlap among top-ranked gene lists — a criterion that rewards methods for agreeing with one another rather than for being right. The most methodologically ambitious attempt to establish a ground truth, Schurch et al. (2016), leveraged 48 biological replicates in yeast to define a reference gene list — yet this reference was itself derived from the same experimental data by subsampling, rendering the evaluation inherently circular. A method that overfits the statistical properties of this specific dataset would appear optimal without being genuinely superior. Froussios et al. (2019) extended benchmarking to the complexity of a eukaryotic genome using *Arabidopsis thaliana*, finding broadly comparable performance across methods — but again without a biologically motivated evaluation criterion. Baik et al. (2020) and Corchete et al. (2020) both relied predominantly on simulated data, with the latter finding comparable performance across multiple methods at various stages of the RNA-seq analysis pipeline. Stupnikov et al. (2021) reported NOISeq as the most robust method and notably found high overlap of GO terms across methods — but crucially, neither the methodology of GO enrichment nor the biological relevance of the recovered terms was discussed in depth, leaving the finding difficult to interpret or build upon. Li et al. (2022) similarly identified multiple top-performing methods using DE gene overlap as the evaluation criterion. Buratin et al. (2023) represents a partial advance, focusing on circular RNA and employing Jaccard similarity — a metric also used in the present study — but remaining confined to a specialised RNA class.

Taken together, the field has been disproportionately reliant on simulated data and, where real data were used, has consistently substituted inter-method agreement for biological validity. The convergence of methods at larger sample sizes, while real, does not resolve this problem — it merely obscures it. In this context, the present study departs from the established paradigm in four substantive ways: (i) we use multiple real datasets and biologically distinct contrasts spanning both mouse and human tissue; (ii) we focus explicitly on the uniqueness rather than the overlap of DE gene sets, treating method-specific genes as potentially informative rather than suspect; (iii) we evaluate biological relevance through detailed GO and Disease Ontology enrichment analysis, with organ-specific interpretation of recovered terms; and (iv) to our knowledge, this is the first benchmarking study to employ Disease Ontology as an evaluation criterion for DE methods.

Our results demonstrate that DELocal consistently identifies a distinct but biologically coherent set of genes across diverse tissues and species — genes that not only differ from those recovered by conventional methods but are enriched for functionally relevant terms that other methods miss entirely. In contrast, the other methods provided largely comparable set of genes among the methods. The differentiating factor for DELocal is its use of neighbourhood concept in identifying differential expression. In practice, this distinction appears to affect highly expressed genes with relatively small changes being ranked higher in the differentially expressed gene lists. We argue that including or excluding the neighbourhood approach in the DE methods is not inherently right or wrong. Rather, DELocal provides a biologically alternative method in the analyses of transcriptomic data. To actually perform the analyses so that the methods can be compared, we provide broadSeq with a robust, extensible framework to conduct this kind of evaluation on any dataset of interest, moving DE method benchmarking from a question of statistical agreement to one of biological insight.

## Methods

### Design and implementation

The broadSeq package is developed for R environment where majority of popular RNA-seq methods for count data are available. The programming style has kept as simple as possible so that any new function can be easily integrated. Majority of plotting functions return ggplot objects to easily construct multi-figure panels. These functions are also return raw data to be used by advanced user for further custom analysis. It has been integrated with ggpubr (Kassambara 2026) for publication-ready figures. All these clustering function returns the subset of SummarizedExperiment data, which is used for final analysis and visualization, in a format which can be further used with any base function.

### RNA-seq Datasets

The raw RNAseq reads of developing mouse organs (Cardoso-Moreira et al. 2019) were first downloaded [Accession E-MTAB-6798] and bad reads were filtered out using FastQC (v. 0.11.8 Andrews 2010), AfterQC (v. 0.9.6 Chen et al. 2017), and Trimmomatic (v. 0.39 Bolger et al. 2014), and ribosomal RNA was removed using Sortmerna (Kopylova et al. 2012). Further the reads were aligned using Salmon (v. 0.99.0 Patro et al. 2017b) to GRCm39 (Ensembl release 104) cDNA.

The Wilms tumor processed dataset were downloaded from the database of Genotypes and Phenotypes (dbGaP) under the accession number phs000471 (Gadd et al. 2017, NCI TARGET Initiative 2017). The dataset is consist of 124 tumor and 6 normal tissue samples.

### Evaluation

The broadSeq [v.1.5.2] provided R packeges limma(v.3.66.0 Ritchie et al. 2015), DESeq2(v.1.50.2 Love et al. 2014), edgeR(v.4.8.2 Chen et al. 2025), EBSeq(v.2.8.0 Leng et al. 2013), NOISeq(v.2.54.0 Tarazona et al. 2011) and DELocal(v.1.11.1 Das Roy et al. 2021) were compared to identify differentially expressed genes whose adjusted p-value or false discovery rate(FDR) were less than 0.05. Thereafter up-regulated genes were ranked on descending order of reported fold change values by each method.

Further the functional profiles of these top 100 ranked genes were compared by clusterProfiler (v.4.18.4 Yu et al. 2012, Xu et al. 2024) either using Gene Ontology (v.3.22.0 Carlson 2025) (biological process) or Disease Ontology (v.4.4.0 Yu et al. 2014). Terms with adjusted p-value less than 0.05 were only reported.

All programs are executed in R version 4.5.2.

The bioconductor R package broadSeq is available at https://bioconductor.org/packages//release/bioc/html/broadSeq.html

## Acknowledgements

We wish to acknowledge the services of the HiLIFE Biodata Unit supported by HiLIFE and CSC—IT Center for Science, Finland, for computational resources.

## Supplementary

**Table S1:**
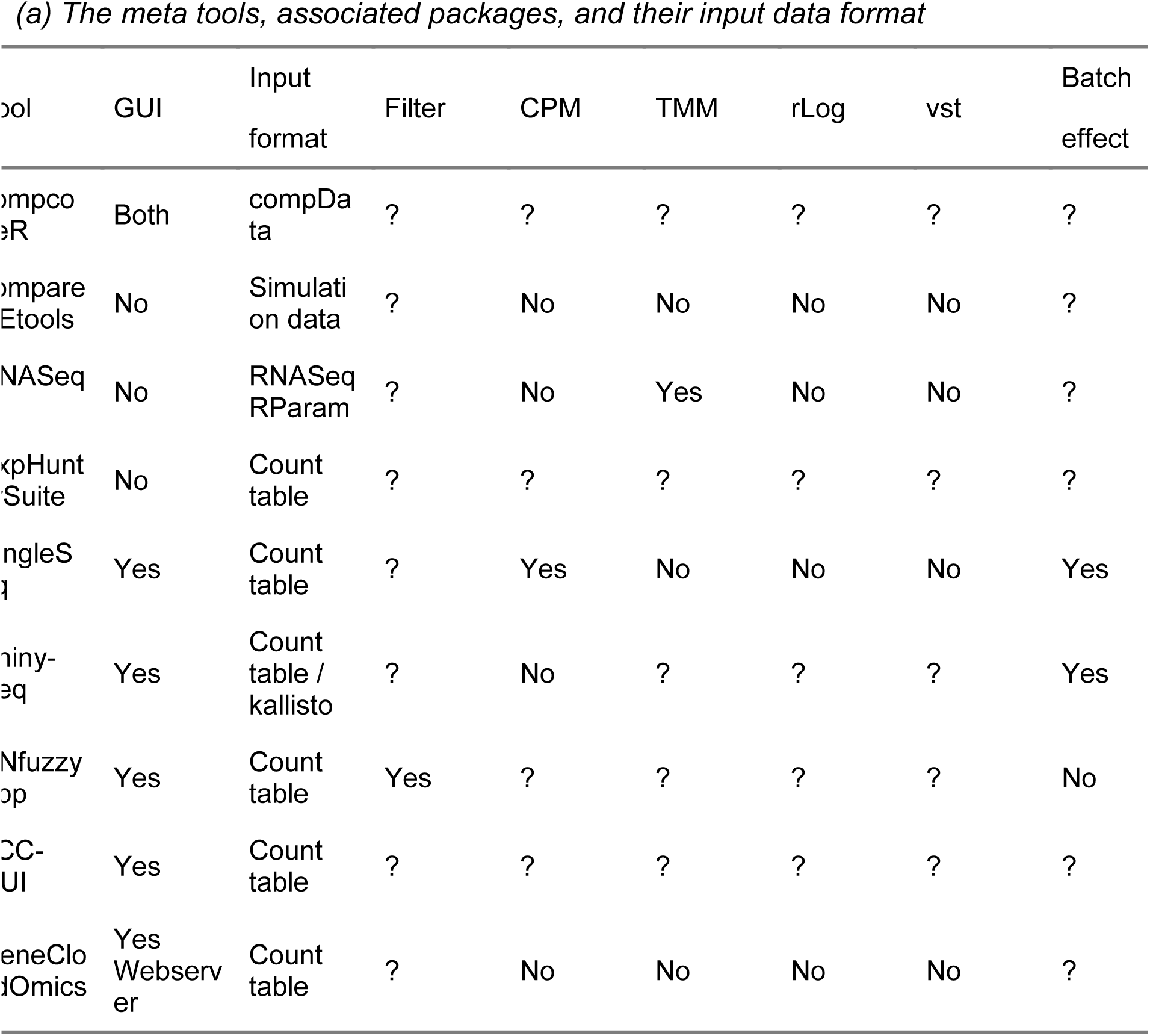
The meta tools and their features.

**Table S2:** The meta tools: Available clustering and differential expression methods.

| Tool | Hierarchical | MDS | PCA | Paired | Group | Multiple methods | compare |
| --- | --- | --- | --- | --- | --- | --- | --- |
| compcoder | No | No | No | ? |  | Yes | Yes |
| compareDEtools | No | No | No | ? |  |  |  |
| RNASeqR | Yes | No | Yes | ? | 2 |  | No |
| ExpHunterSuite | ? | ? | ? | ? |  | Yes | No |
| BingleSeq | No | No | Yes | ? |  | Yes | Yes |
| Shiny-Seq | Yes | No | Yes | ? |  | No | No |
| RNfuzzyApp | Yes | No | Yes | Yes |  | Yes | No |
| TCC-GUI | Yes | Yes | Yes | ? | 3 | Yes | No |
| GeneCloudOmics | Yes | No | Yes | ? |  | Yes | No |

**Figure S1:**
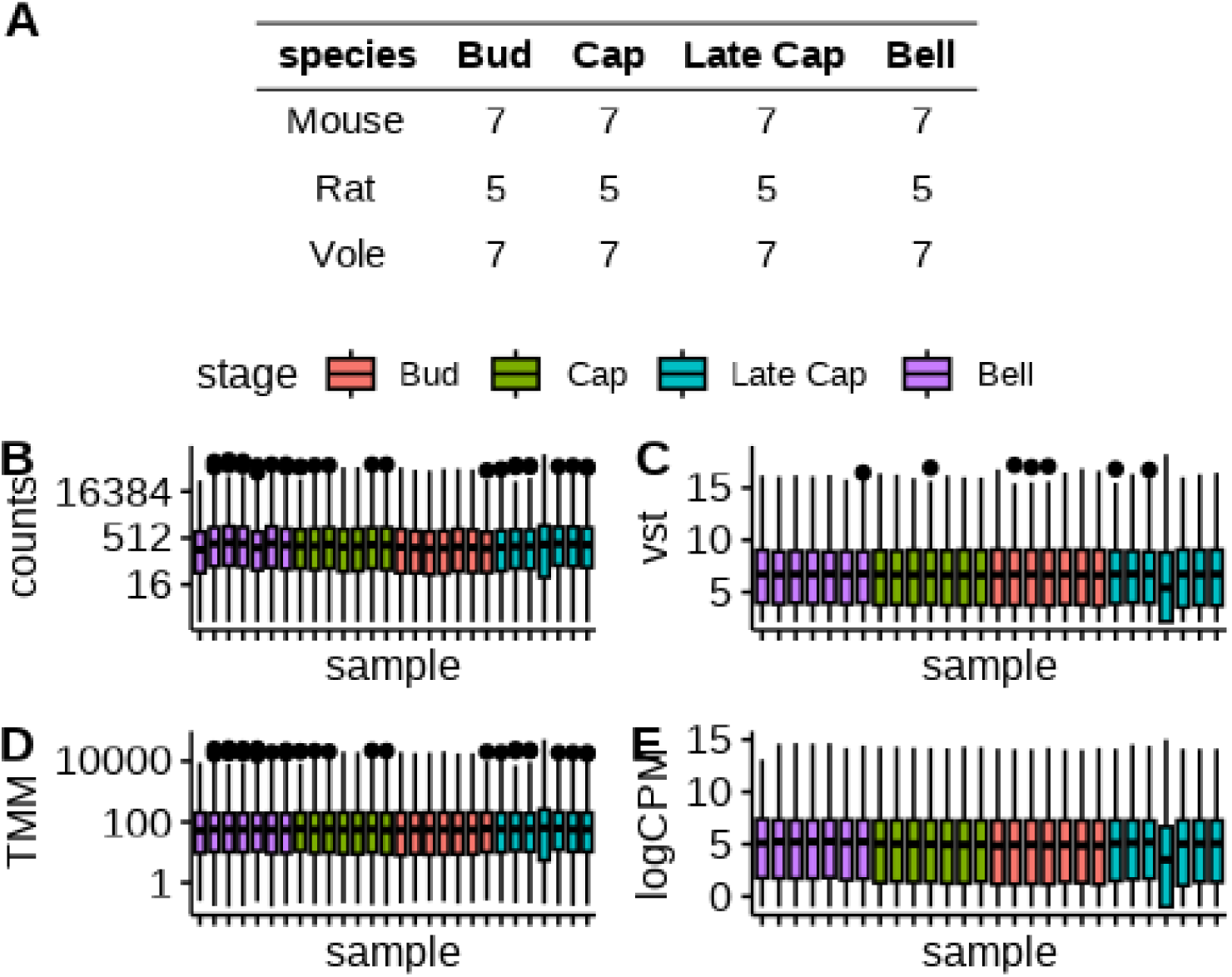
The broadSeq (A) Data used for to demonstrate visualization. (B-E) Normalization and Transformation.

**Figure S2:**
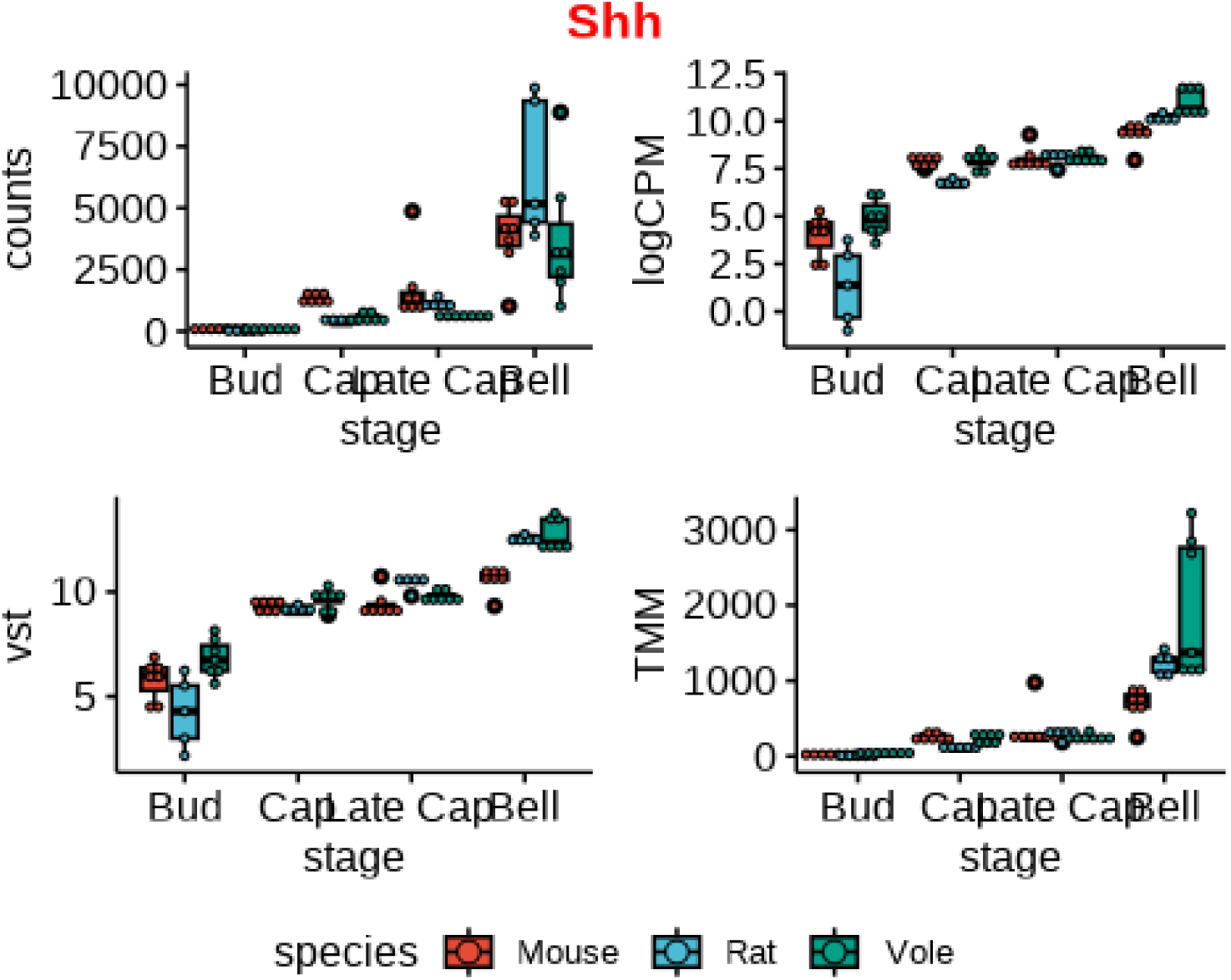
Easy exploration of gene expression across different normalization.

**Figure S3:**
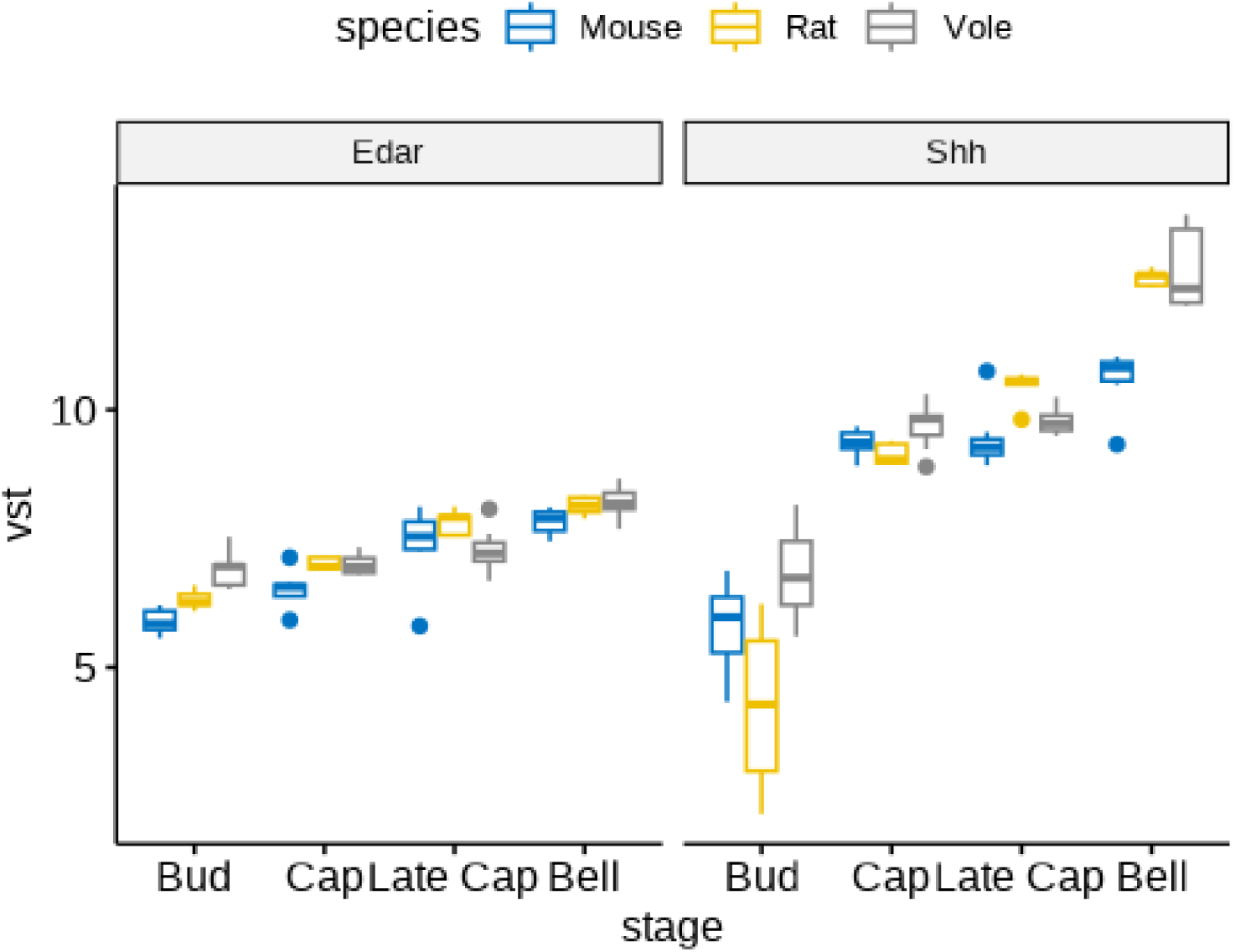
Exploration of multiple gene expression.

**Figure S4:**
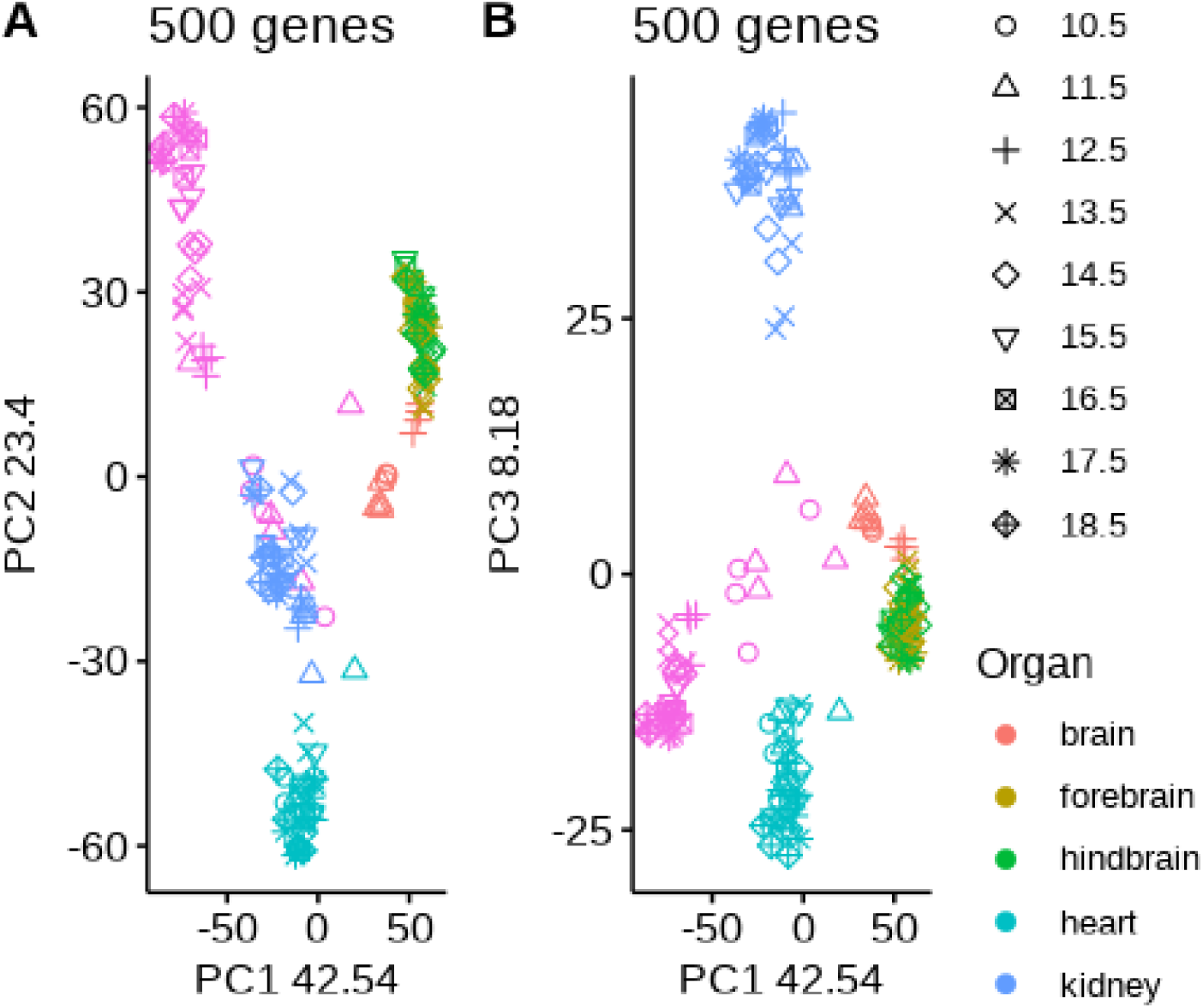
Principal components 1, 2 and 3 of mouse developmental dataset are driven by variations of genes specific to heart, liver and kidney and fails to capture developmental variation

**Figure S5:**
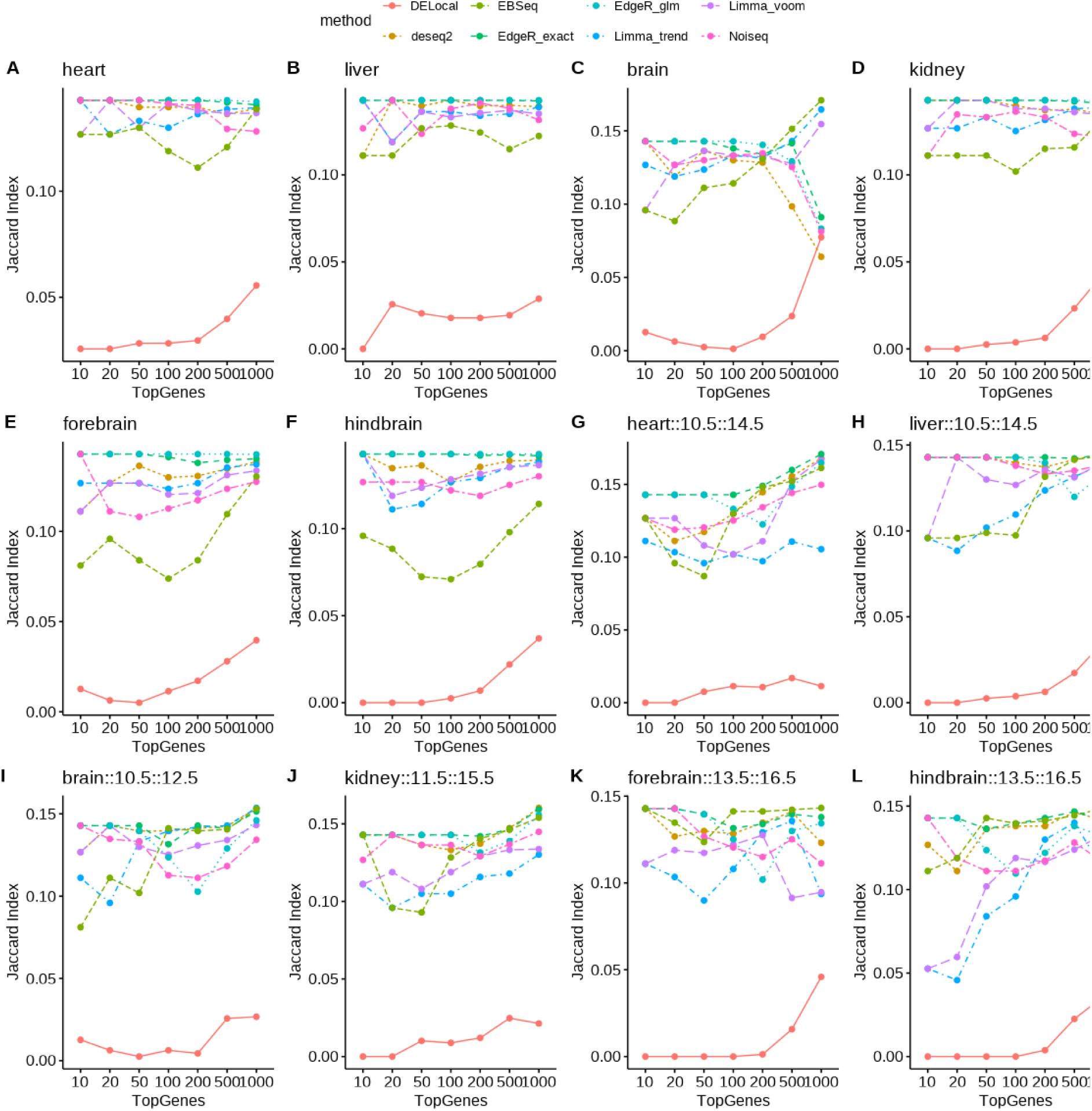
For each contrast and different number of top genes jaccard index were measured between methods. DELocal shows least similarity with rest of the methods.(A-F) Organ specific up-regulated genes. (G-L) Developmental up-regulated genes in a organ (early and mid developmental timepoints are shown in the figure titles)

**Figure S6:**
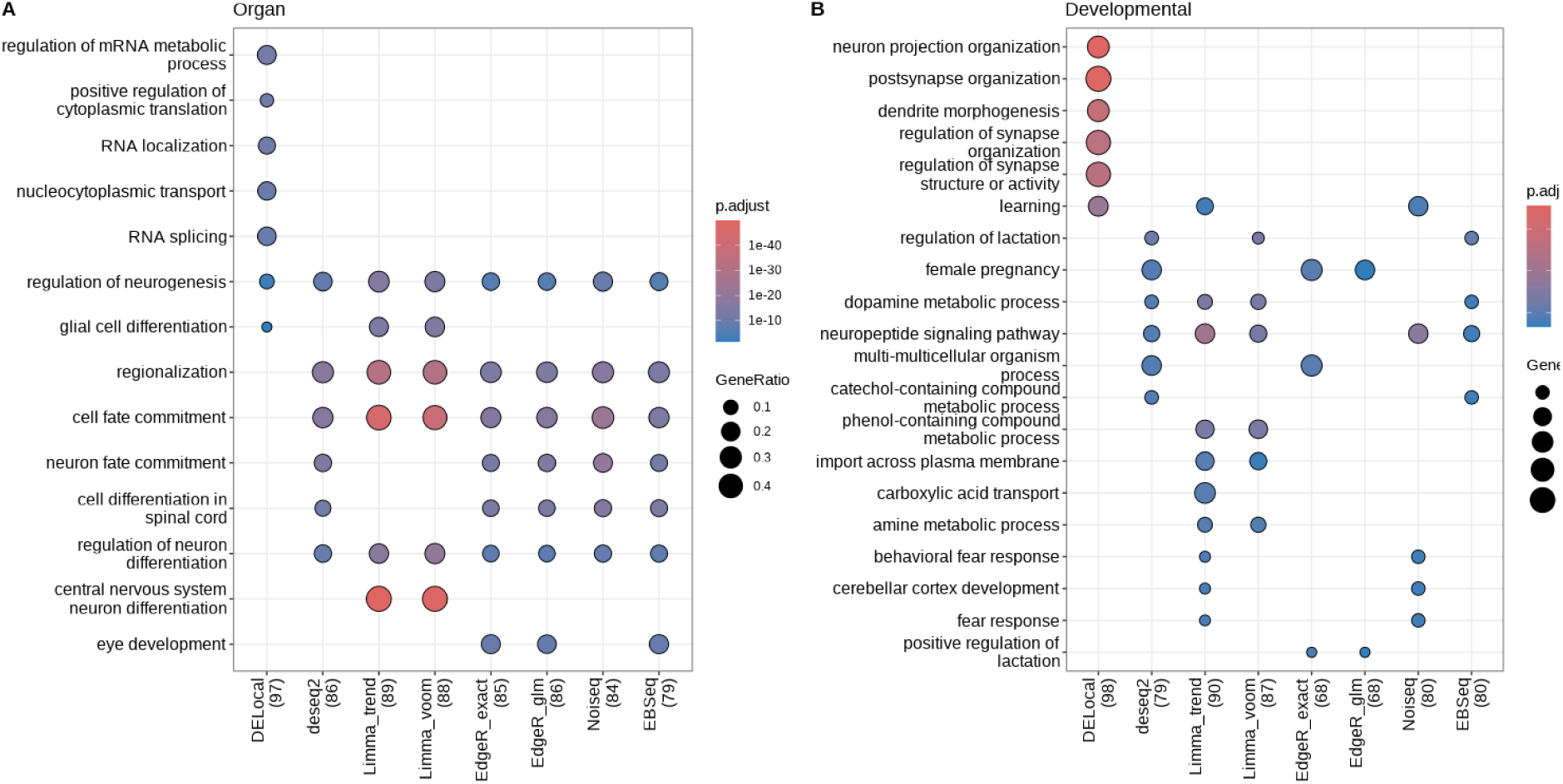
Brain (A) Organ identity (B) Developmental trajectory.

**Figure S7:**
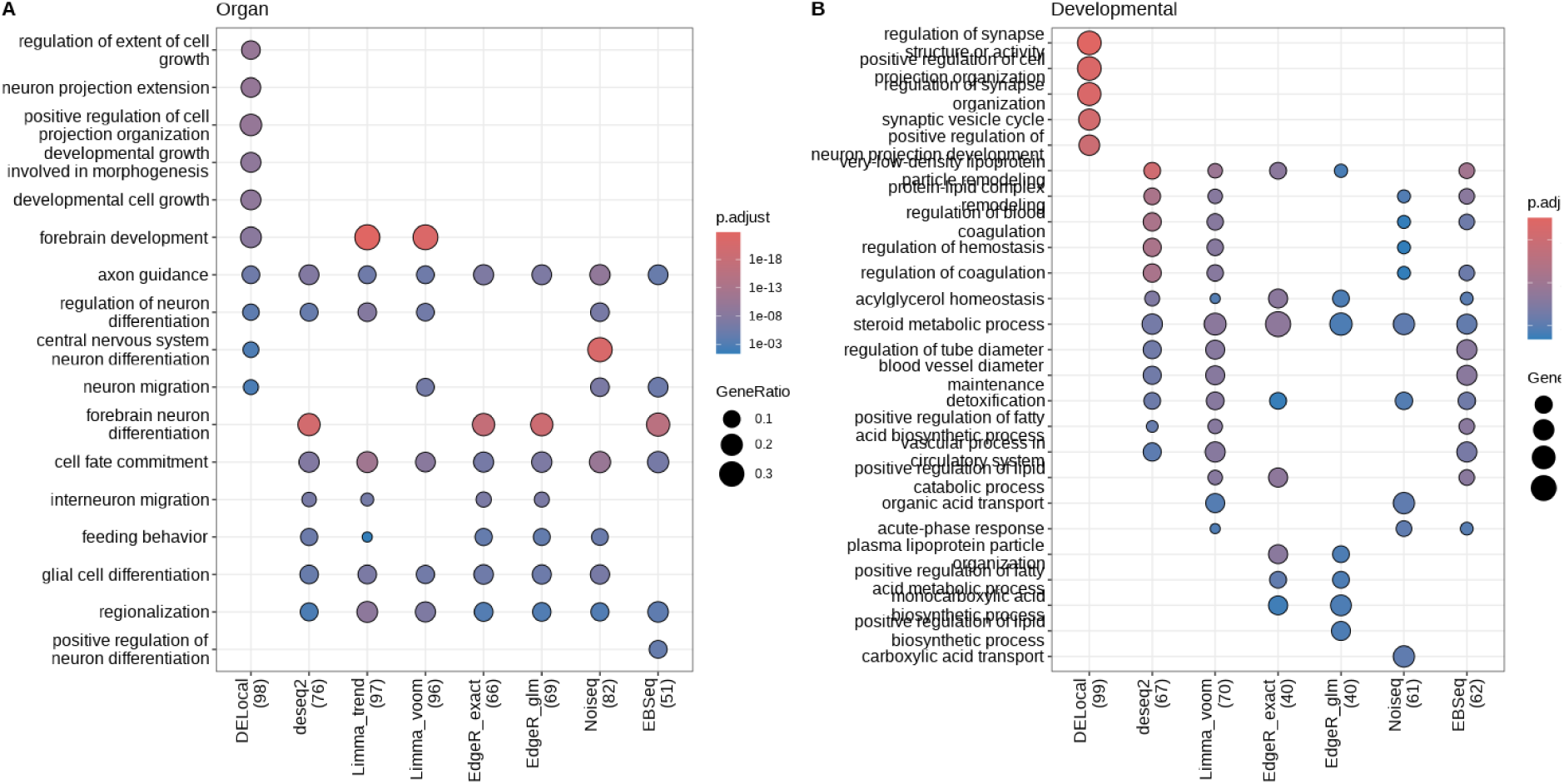
Forebrain (A) Organ identity (B) Developmental trajectory.

**Figure S8:**
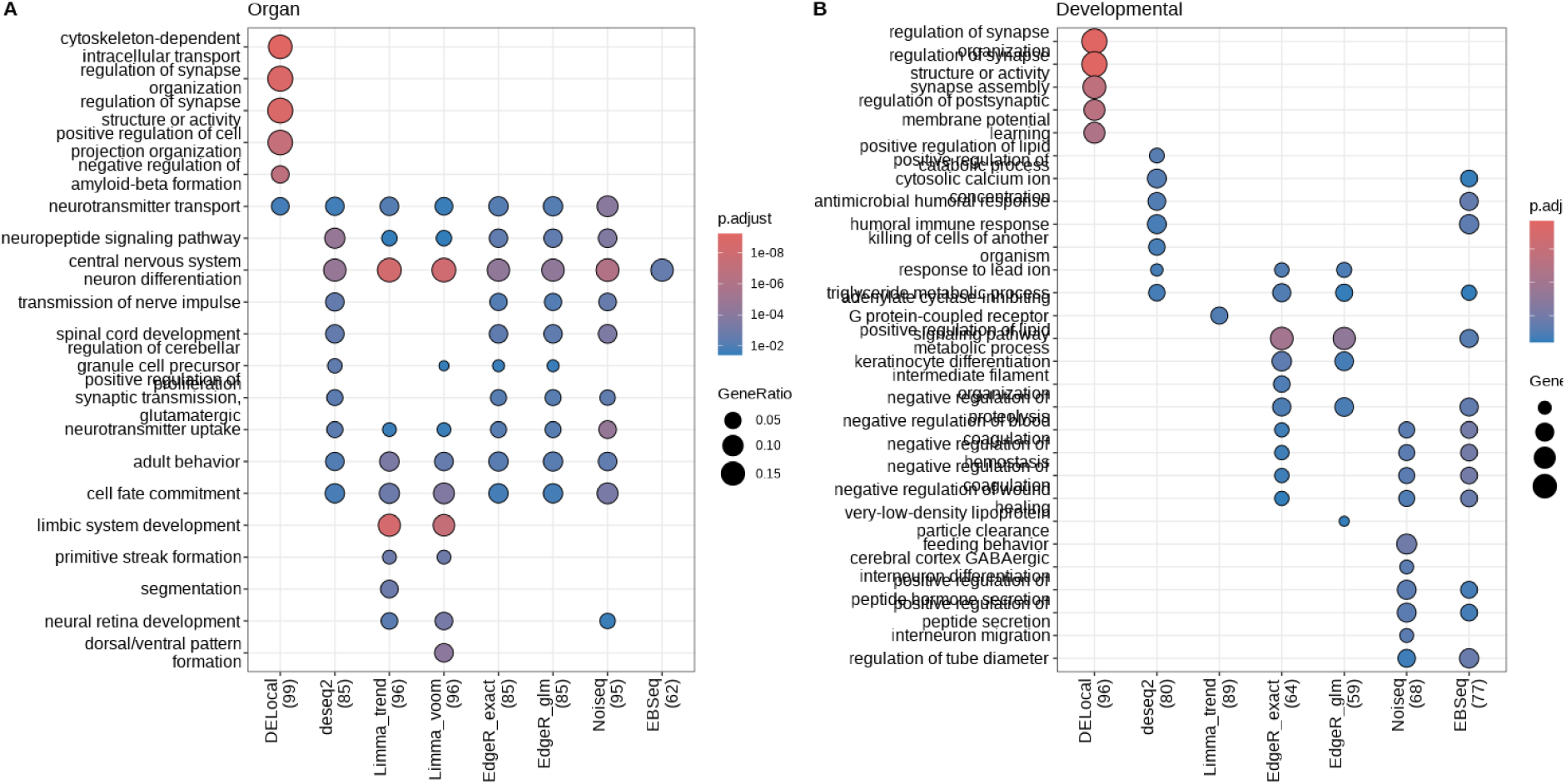
Hindbrain (A) Organ identity (B) Developmental trajectory.

**Figure S9:**
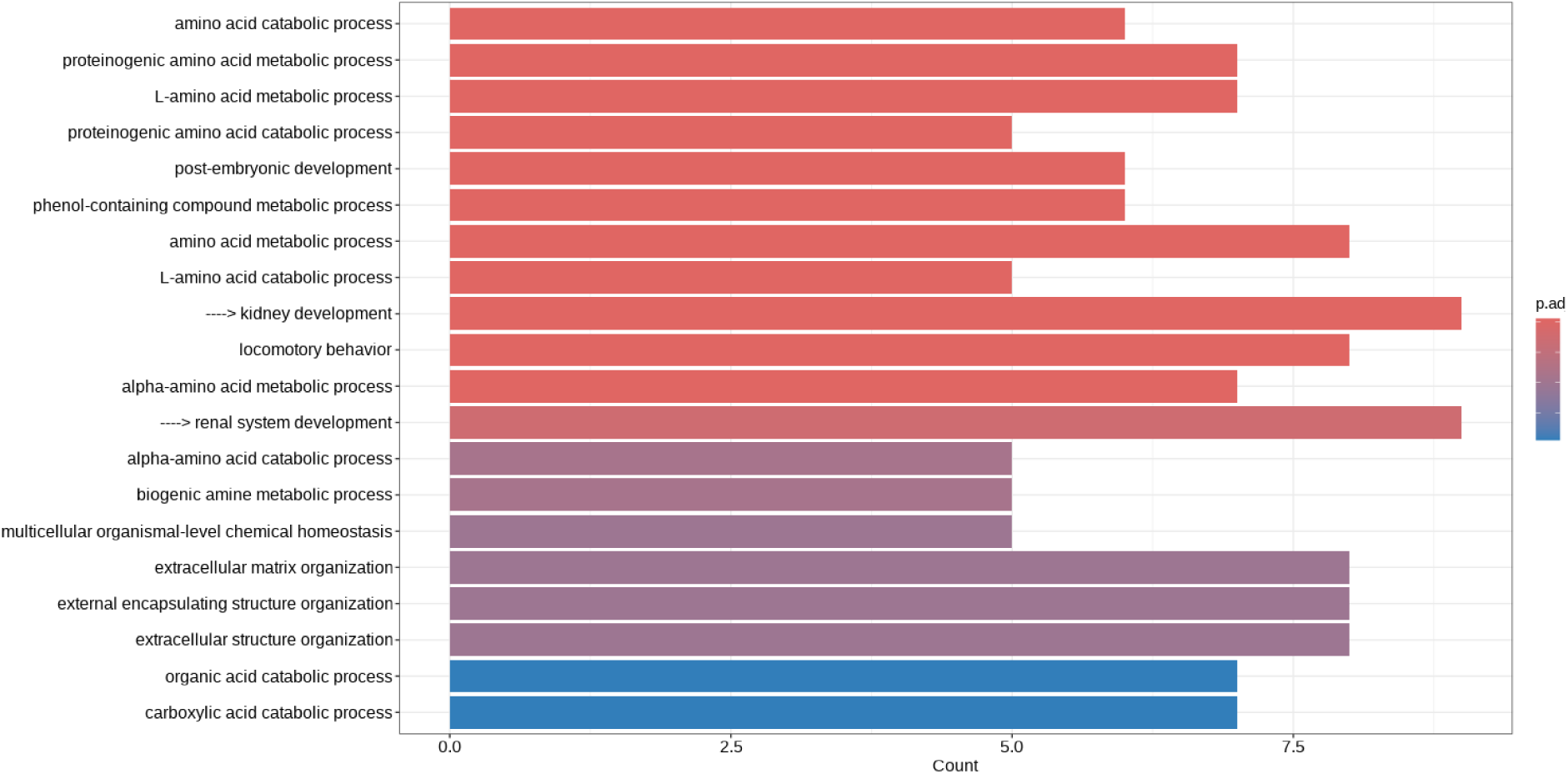
Top 20 significant GO terms from DELocal’s top 100 gene list includes kidney related GO terms.

**Figure S10:**
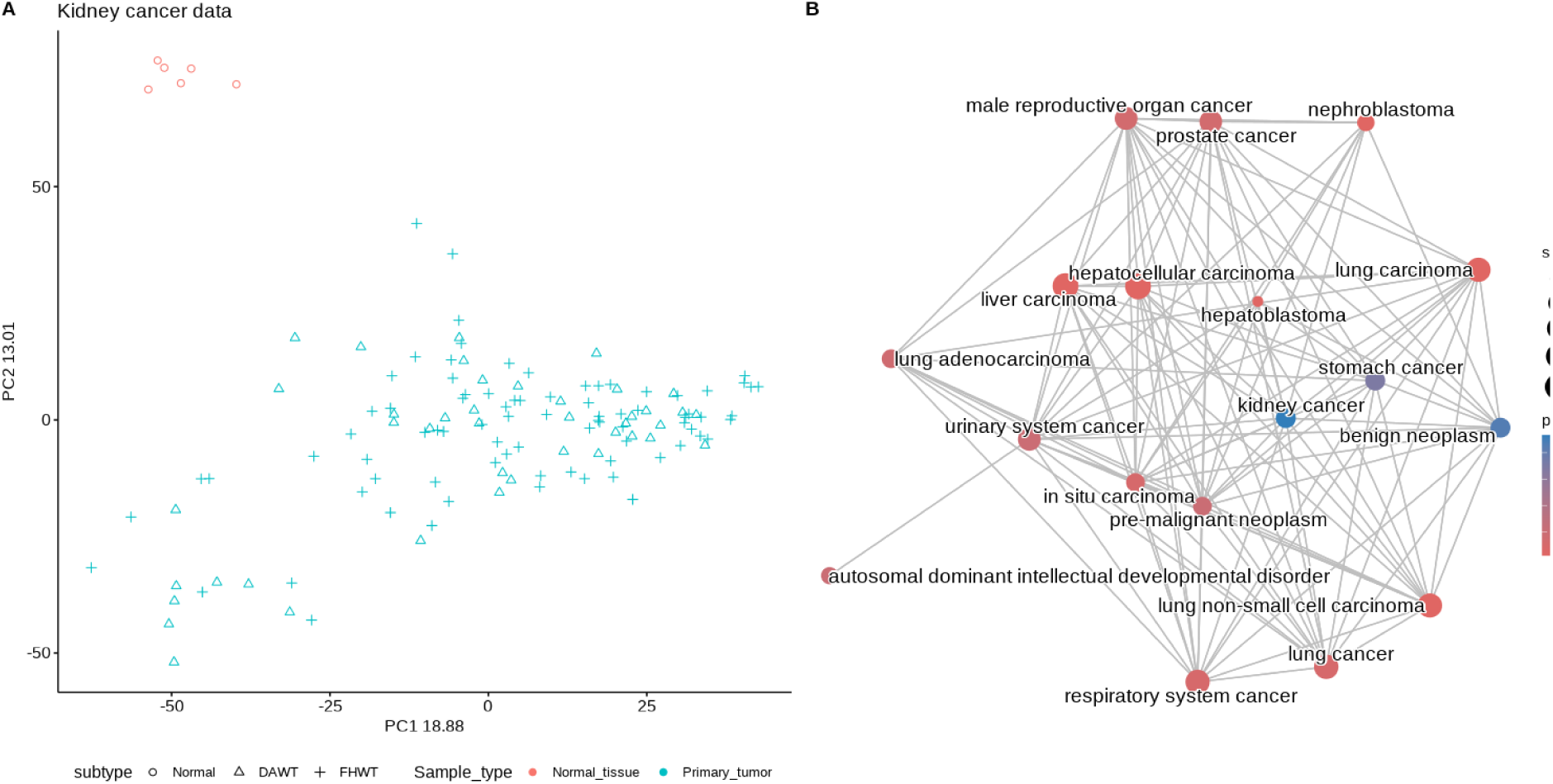
Kidney cancer data. (A) PCA plot of shows clear separation normal tissue type from primary tumor types. (B) DELocal identified genes also includes ‘kidney cancer’ term.

**Figure S11:**
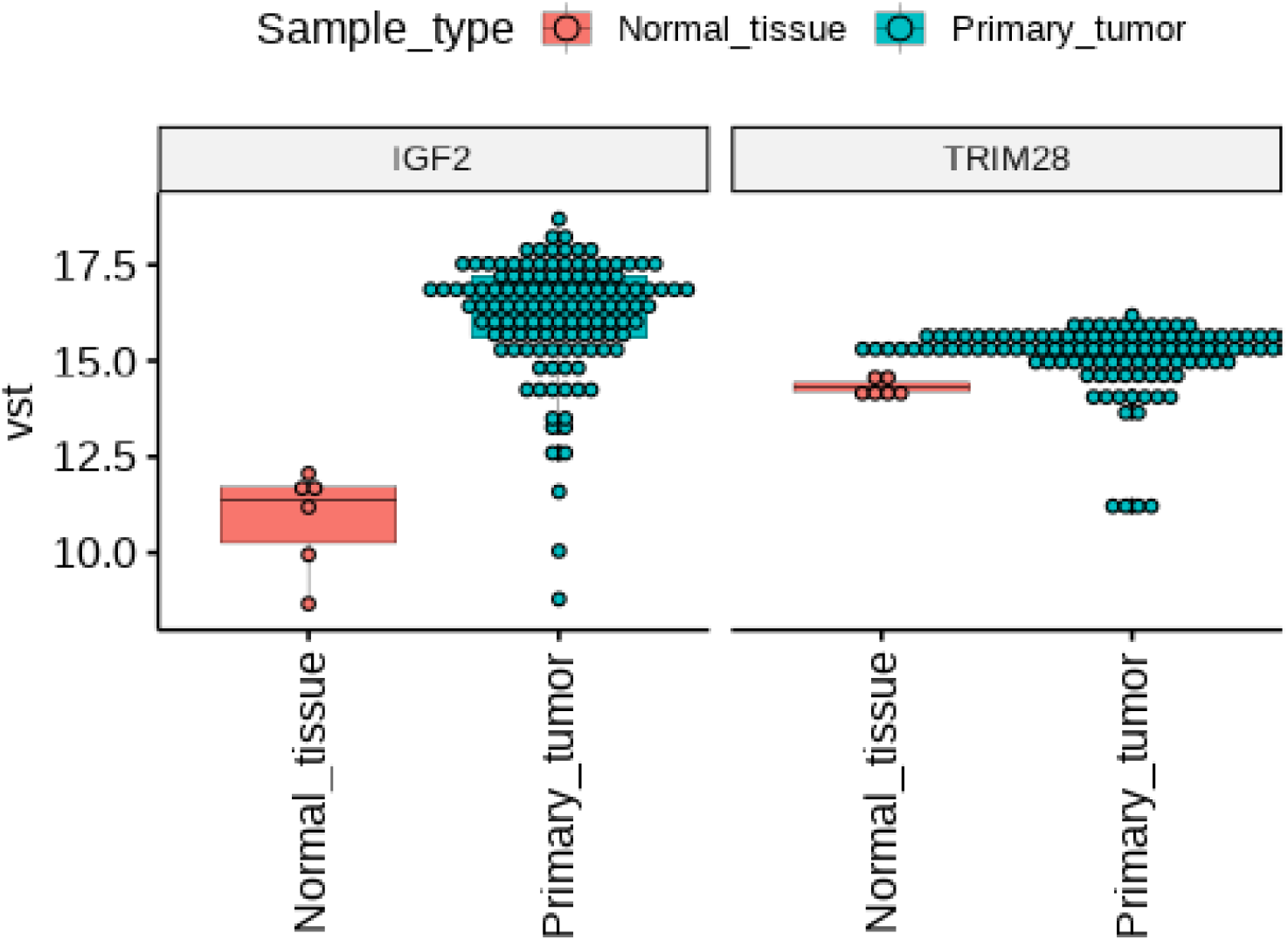
Normalised expression measured through VST function of DESeq2 of IGF2 and TRIM28.

